# pH-Triggered Assembly of Endomembrane Multicompartments in Synthetic Cells

**DOI:** 10.1101/2021.08.25.457616

**Authors:** Félix Lussier, Martin Schröter, Nicolas J. Diercks, Kevin Jahnke, Cornelia Weber, Christoph Frey, Ilia Platzman, Joachim P. Spatz

**Affiliations:** Department of Cellular Biophysics, Max Planck Institute for Medical Research, Jahnstraße 29, D- 69120 Heidelberg, Germany; Institute for Molecular Systems Engineering (IMSE), Heidelberg University, Im Neuenheimer Feld 225, D-69120 Heidelberg, Germany; Max Planck Institute for Medical Research, Biophysical Engineering Group, Jahnstraße 29, D-69120 Heidelberg, Germany; Department of Physics and Astronomy, Heidelberg University, D-69120 Heidelberg, Germany; Max Planck School Matter to Life, Jahnstraße 29, D-69120 Heidelberg, Germany

## Abstract

Bottom-up synthetic biology thrives to reconstruct basic cellular processes into a minimalist cellular replica to foster their investigation in greater details with a reduced number of variables. Among these cellular features, the endomembrane system is an important aspect of cells which is at the origin of many of their functions. Still, the reconstruction of these inner compartments within a lipid-based vesicle remains challenging and poorly controlled. Herein, we report the use of pH as external trigger to self-assemble compartmentalized giant unilamellar vesicles (GUVs) by either bulk, or droplet-based microfluidics. By co-encapsulating pH sensitive small unilamellar vesicles (SUVs), negatively charged SUVs and/or proteins, we show that acidification of the droplets efficiently produces GUVs while sequestrating the co-encapsulated material with flexibility and robustness. The method enables the simultaneous reconstruction of more than a single cellular phenotype from the bottom-up, corresponding to an important advancement in the current status quo of bottom-up synthetic biology.

## Introduction

The idea to fully reconstruct eukaryotic cells in a laboratory is, at first glance, far-fetched due to all the requirements that must be fulfilled in order to build life from scratch. Hence, achieving the construction of a living system from non-living building blocks would tremendously impact various aspects of cellular biology, ranging from revolutionizing our understanding of the origin of life^1^, to the production of man-made artificial cells to fight cancer.^2^ Toward this aim, synthetic biology dissects, isolates and reconstructs cellular processes through the assembly of well-characterized molecular building blocks. This holistic vision is thus promoting and improving our current understanding of individual cellular components, but also fostering our ability to investigate their collective and emerging properties. Up to now, various cellular components, or key functionalities of a living cell such as energy production^3, 4^, metabolism^5, 6^, signaling^7^, protein expression^8, 9^, growth^10^, division^11^ and cytoskeleton^4, 12^ have all been individually reconstructed within cell-sized systems and allowed their investigation in greater detail. To perform a majority of these tasks, eukaryotes rely on a vast and complex endomembrane system which segregate various cellular functions into specialized compartments referred to as organelles. The presence of organelles, and hence the concept of compartmentalization, enables the co-existence of chemically distinct reactions in spatially confined reactors while allowing multi-step reactions, and sustaining chemical gradients. Thus, achieving the controllable genesis of compartments within synthetic eukaryotes still consists of an important leap in bottom-up synthetic biology.

In fact, compartments have already been reconstructed within lipid-based vesicles, often referred to as vesosomes, and have raised attention in the field of drug delivery by minimizing passive leakage of therapeutic drugs.^13^ However, vesosome assembly is achieved through bulk processes, where the lipid composition, polydispersity, throughput, and reproducibility are limited.^13-15^ These methods are thus not easily translated for the controlled bottom-up reconstruction of a synthetic eukaryote for synthetic biology applications. Most of these limitations may be circumvented through the usage of droplet-based microfluidics to improve precision and manipulation. For instance, microfluidic technology can be implemented to generate water-in-oil-in-water (W/O/W) double emulsions and allows for encapsulation of small^10, 16, 17^ or large lipid compartments^18^ inside GUVs. However, W/O/W droplets applied to bottom-up synthetic biology have shown only the use of an *a priori* limited set of lipids (i.e. mostly phosphatidylcholine (PC)-based lipids). Moreover, this approach often relies on the use of non-ionic surfactants to foster the spontaneous dewetting of the excess of oil by minimizing the total interfacial energy^19, 20^ and avoid membrane defects.^21, 22^ In addition, fabrication and passivation of microfluidic devices for W/O/W droplet production remains challenging. Taken together, novel and simple methods to effectively mimic the complex endomembrane system in synthetic cell, while enabling a broad and diverse lipid composition would greatly benefit the field of bottom-up synthetic biology.

Toward this aim, Göpfrich and colleagues have recently reported the self-assembly of multicompartment vesicles within water-in-oil droplets, referred to as droplet-stabilized GUVs (dsGUVs), by co-encapsulating SUVs of opposite charges.^23^ In this study, the droplets were stabilized by a mixture of uncharged PEG-based fluorosurfactant and a negatively-charged PFPE carboxylic acid fluorosurfactant (namely Krytox). A net negative charge at the W/O droplet interface initiated the selective recruitment and fusion of the cationic SUVs at the droplet periphery. The resulting dsGUVs sequestered the negative SUVs, which remained within the vesicle lumen. Albeit promising, the need of cationic lipids (i.e. DOTAP) to recreate an endomembrane system in GUVs still lacks flexibility, is intricately unnatural and highly cytotoxic.^24, 25^ As an alternative to permanent cationic lipids, pH sensitive lipids bearing chemical functional groups capable to modulate their ionic state as a function of pH could circumvent this problem.^26^

Herein, we present the pH-mediated reconstruction of an endomembrane system within GUVs through a W/O emulsion using both, bulk and microfluidic approaches. Importantly, besides the ability to encapsulate different compartments, the use of pH to trigger the charge-mediated assembly of dsGUVs greatly improves the production efficiency of free-standing GUVs, a major hurdle in the droplet-stabilized method. In addition, and as a proof-of-concept, the method facilitated the reconstruction of a F-actin network, with and without an endomembrane system. These results showcase the potential of the pH-triggered assembly of GUVs to reconstruct more than a single cellular component, an important leap in bottom-up synthetic biology, where the complexity of the synthetic eukaryote can be incremented.

## Results

### pH-mediated assembly of dsGUVs

As a first step, we investigated the potential use of pH to mediate the assembly of dsGUVs in bulk, by combining and vortexing the water and oil phases for rapid prototyping of the experimental conditions. Towards this end, SUVs containing the pH sensitive lipid *N*-(4-carboxybenzyl)-*N,N*-dimethyl-2,3-bis(oleoyloxy)propan-1-aminium (DOBAQ) were suspended in citrate buffer at various pH and encapsulated within W/O droplets stabilized by an oil-surfactant mixture composed of 2.5 mM PEG-based fluorosurfactant and 10 mM Krytox in HFE-7500. Imaging by confocal laser scanning microscopy (CLSM) revealed that at pH 6, the fluorescence signal associated to Lissamine Rhodamine B (Liss Rhod B)-labelled lipids supplemented to the SUVs was uniformly distributed within the droplets’ lumen (Figure 1A). Upon reduction of the intraluminal pH, we observed an increased recruitment of the pH sensitive SUVs to the interface of the W/O droplets. A complete recruitment and fusion of the encapsulated SUVs at pH 5 led to the assembly of droplet-stabilized GUVs (Figure 1A). FRAP measurements confirmed the successful assembly of a supported lipid bilayer. The measured diffusion coefficient (2.99 ± 0.34 µm^2^/s) of Liss Rhod B-labelled DOPE lipids matches similar values reported for dsGUVs (Figure 1B).^4, 23^

**Figure 1:**
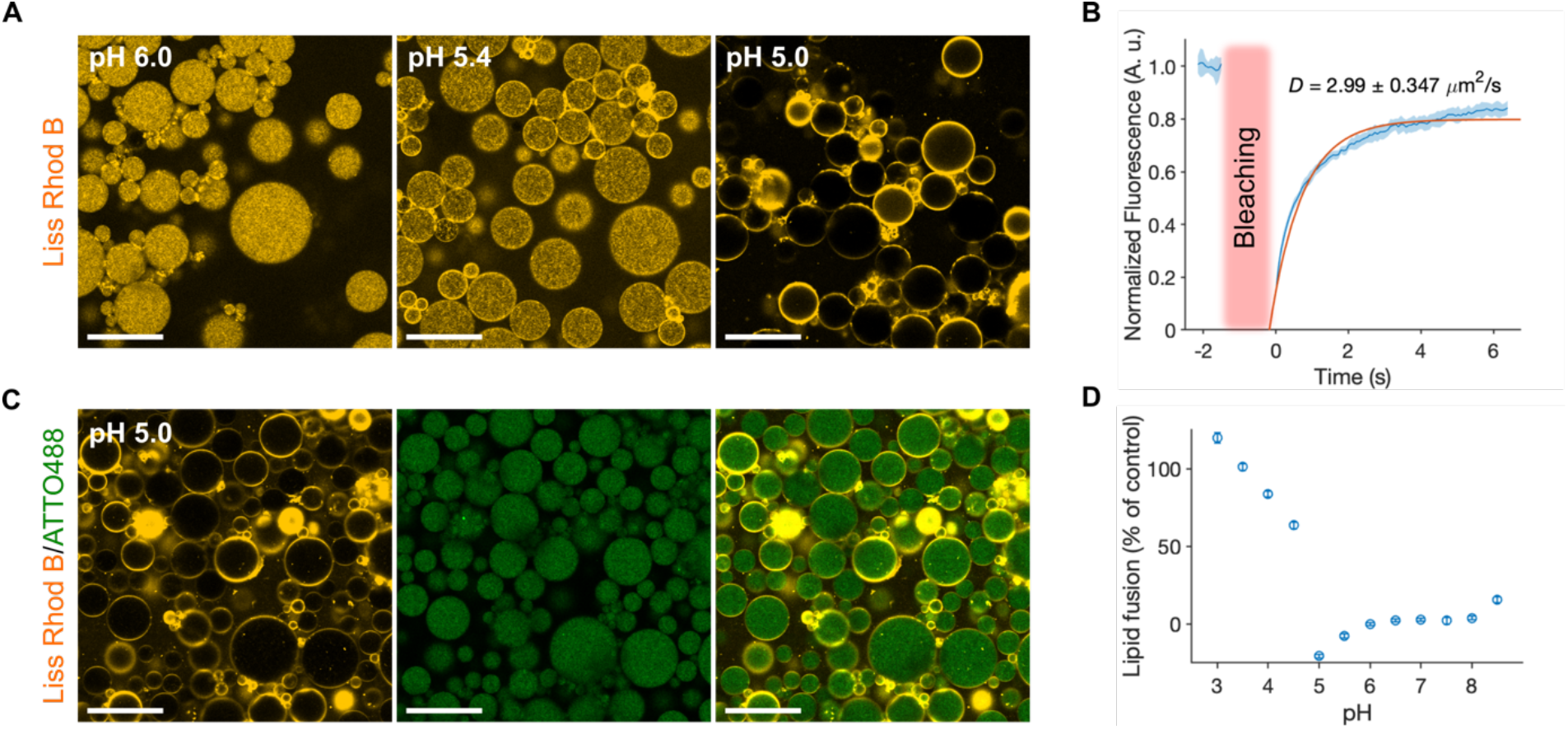
pH-mediated assembly of dsGUVs. (**A**) Representative confocal images of the encapsulated pH sensitive SUVs composed of DOBAQ/eggPG/eggPC/Liss Rhod B-labelled DOPE (60/20/19.5/0.5 mol%) within the W/O droplets stabilized by 1.4 wt/wt% PEG-based fluorosurfactant and 10 mM Krytox. Various 50 mM citrate buffers we used to adjust the pH of the water phase. Scale bars: 50 μm. (**B**) Fluorescence recovery after photobleaching (FRAP) of dsGUVs assemble through a pH trigger. Mean ± S.D. (*n = 11*) are presented. The bleached area is highlighted with the red rectangle. The mean-normalized fluorescence intensity values within the circular bleached area (2.5 µm radius) are plotted as a function of time. The orange line represents an exponential fit (*R*^2^ = 0.9301), which was further used to extract the diffusion coefficient *D* of the lipids in the dsGUVs. The extracted value of *D* was 2.99 ± 0.34 µm^2^/s, similar to previously reported value for dsGUVs thus confirming the formation of a supported-lipid bilayer within the W/O droplet. (**C**) Self-assembly of multicompartment dsGUVs in presence of 50 mM citrate buffer at pH 5. The pH sensitive SUVs were co-encapsulated with negatively charged SUVs composed of DOPC/DOPG/ATTO488-labelled DOPE (79.5/20/0.5 mol%). Scale bars: 50 μm. (**D**) FRET experiment measuring lipid mixing of pH sensitive SUVs. SUVs composed of DOBAQ/eggPG/eggPC/Liss Rhod B-labelled DOPE/NBD-labelled DOPE (60/20/18/1/1 mol%) were mixed with unlabeled negatively charged SUVs at various pH. At a pH below 5.0, a significant mixing of the SUVs was observed, depicted by the abrupt rise in fluorescence intensity resulting from the unquenching of the NBD reporter by Liss Rhod B. Mean ± S.D. are presented (*n = 3)*.

To characterize the fusion behavior of DOBAQ-SUVs as a function of pH at the droplet interface, we first assessed the pK_a_ of the DOBAQ lipid in SUVs spectroscopically throughout a 2-(*p*-toluidino)-6-naphthalene sulfonic acid (TNS)-based assay. TNS is a fluorescent reporter whose fluorescence is attenuated in a hydrophilic environment and widely applied to evaluate the pK*a* of lipid-based nanoparticles.^26-29^ Due to its intricate negative charge (Figure S1), TNS is more readily attracted to positively charged lipid membranes. The resulting increase in lipophilicity of the local vicinity unquenches the fluorescence of the TNS probe. Thus, the fluorescence of TNS serves as an indicator of the surface charge of lipid-like vesicles. By varying the pH of the SUVs suspension in presence of TNS, we evaluated the surface charge of DOBAQ, and hence it’s pK_a_. Through the TNS-assay (see Supplementary Information), the pK_a_ of DOBAQ was estimated to be 4.35 by fitting a sigmoid function (Figure S2), and by evaluating the pK_a_ as the point at half-maximum, where 50% of the DOBAQ would be protonated. This value is in good agreements with zeta potential measurement^30^ and fusion assays as a function of pH reported elsewhere.^31^

Up to now, the charge-mediated generation of multicompartment dsGUVs was limited to either the use of cationic SUVs containing DOTAP lipids which were preferentially recruited at the negatively charged droplet periphery while entrapping negatively charged SUVs,^23^ or via the encapsulation of an excess of negatively charged SUVs in presence of Mg^2+^ ions.^32^ Here, we investigated the use of pH to trigger the assembly of multicompartment dsGUVs from the bottom-up, and avoid the use of permanently cationic lipids. To this extent, two SUV populations were suspended in various citrate buffers containing no Mg^2+^ ions and co-encapsulated within W/O droplets stabilized by the same oil-surfactant mixture (Figure 1C). Here, one population of SUVs were pH sensitive (i.e. DOBAQ/eggPG/eggPC/Liss Rhod B-labelled DOPE) while the other SUVs possessed no pH sensitive motif and were negatively charged at pH 5 and 7.4 (i.e. DOPC/DOPG/ATTO488-labelled DOPE) (Figure S2). Upon emulsification at pH 5, we observed the selective recruitment and fusion of the pH sensitive SUVs on the droplet periphery (Figure 1C), while limited to no recruitment was observed at higher pH (Figure S3). Interestingly and importantly, no lipid mixing, (i.e. fusion) in between the two SUV populations was detected within the droplets as the fluorescent signal associated to the pH sensitive SUVs was solely detected at the droplet periphery at pH 5 (Figure 1C, Fig S3).

FRET measurements were applied to further understand the preferential fusion of pH sensitive SUVs to the droplet periphery over fusion with negatively charged SUVs. Towards this end, using the well-established NBD-Liss Rhod B FRET pair, we measured the lipid mixing between the pH sensitive SUVs and the negatively charged SUVs as a function of pH (Figure 1D). As the pH decreased, a negligible lipid mixing was observed between the SUVs at pH > 5, while an abrupt rise in fluorescence signal was detected at lower pH. This increase in fluorescence signal was associated to the unquenching of the NBD-labelled lipid upon mixing. Interestingly, we detected the lowest lipid mixing exactly at pH 5, corroborating the minimal interaction in between SUVs population encounter at pH 5 and which then preferentially fuse the pH sensitive SUVs to the droplet periphery. This low interaction is an improvement compared to the previously reported assembly of multicompartment dsGUVs employing cationic SUVs in absence of Mg^2+^ ions.^23^

To compare both system, we generated the multicompartment dsGUVs by mixing cationic and negatively charged SUVs in absence of Mg^2+^ at pH 7.4 (Figure S4A).^23^ When produced, cationic SUVs were preferentially recruited at the droplet interface, but also presented a homogeneous fluorescence signal within the droplet’s lumen (Figure S4A). This signal originated from partial lipid mixing between the cationic and negatively charged SUVs, as confirmed by the FRET assay (Figure S4B). Hence, usage of pH sensitive SUVs rather than permanent cationic lipids to assemble multicompartment dsGUVs corresponds to an improved method due to the minimal lipid mixing in between compartments. Moreover, the pH sensitive lipids enable the formation of multicompartment free standing GUVs (see Section: Usage of pH improve the SUVs to GUVs conversion) possessing a net negative charge -30 ± 1 mV (*n =* 3) in physiological conditions as measured by ς-potential (Figure S2C). This is an important prerequisite for further investigation and usage of GUVs for *in vitro* studies since cationic lipids are highly potent toward cellular membrane.^24, 25^

### Engineering of compartmentalized dsGUVs assembly in bulk and microfluidics

To mimic the physical confinement of cells, a key aspect of bottom-up synthetic biology resides in the capability to engineer relatively uniform cell-sized compartments. To generate uniform droplets in a high throughput manner, various droplet-based microfluidics technologies are routinely used in order to generate emulsions directly applied for synthetic biology.^5, 6, 20, 33-35^ Methods enabling a rapid prototyping of various experimental conditions in bulk while being directly translated to a microfluidic platform offer great advantages. To this extend, the charge-mediated assembly of GUVs using W/O droplets possesses such an unique feature. In this approach the droplet size dictates the size of the final vesicle, which can be mitigated, to some extent, by the flow rates of the continuous and dispersed phase, or through the use of different channel geometries. Thus, we evaluated if the use of pH could impede such rapid translation toward microfluidics.

By co-encapsulating pH sensitive and negative SUVs within W/O droplets at pH 5, the assembly of compartmentalized dsGUVs in the average size of 40 µm was achieved by implementation of double aqueous inlet microfluidic device (Figure S5 A-C; Supplementary video S1). However, we observed temporal variation in pressures of the inlets due to aggregation between SUVs upon exposition to the citrate buffer prior their encapsulation in W/O droplets, thus rendering the translation challenging and slightly affecting the homogeneity of the assembled droplets (Figure S5D).

To assemble small cell-sized compartments, we investigated the use of microfluidic devices possessing a smaller channel width and investigate the translation capability of the methods to various microfluidic module. Hence, we applied the pH-mediated approach to assemble compartmentalized dsGUVs employing a mechanical splitter, possessing channels of 2 µm as smallest feature (Figure S6A-B).^30^ With such small channels, we further observed a rapid clogging of the microfluidic channels upon mixing SUVs and citrate buffer which resulted in poor homogeneity, and high polydispersity compared to emulsification by shaking (Figure S6C-D). Consequently, the use of a low pH buffer resulted in a limited translation of this method to microfluidic platforms possessing small channel geometry, and could not be deployed universally.

Since the generation of dsGUVs relies on the generation of an emulsion, we exploited the capacity of the continuous oil phase to externally manipulate the pH of W/O droplets.^36, 37^ Consequently, the acidification of the W/O droplets could be externally controlled after the co-encapsulation of the two SUVs population and splitting of the droplets to prevent potential clogging of the small channels. To first evaluate this hypothesis, we produced dsGUVs encapsulating pH sensitive SUVs in a well-buffered aqueous solution composed of 50 mM KH_2_PO_4_/K_2_HPO_4_ pH 7.4 and generated W/O droplets by the shaking method. For these experiments the oil-surfactant mixture was supplemented with various concentration of acetic acid, a small organic acid soluble in both the aqueous and the fluorinated oil phase. The shaking method allowed a rapid prototyping of various lipid compositions, buffers and surfactant-oil mixture with minimal volume of constituents rather than directly applying microfluidics. In all cases, we kept a water to oil ratio of 1:2, with typical volumes of 10:20 μL respectively. Upon increasing the concentration of acetic acid up to 36 mM, we observed a significant recruitment and fusion of the pH sensitive SUVs to the droplet periphery, while none or negligible recruitment was observed at lower concentration of acetic acid with these lipids and buffer composition (Figure 2A). Interestingly, by increasing the acid concentration, we observed a reduction of the droplet size. This was rationalized by a reduction of the interfacial tension which favors the breaking up of droplets due to the increase in ionic strength within the W/O droplets by the acid. This increase in ionic strength also promotes the concomitant adsorption of further ionic surfactant (i.e. Krytox) at the interface, which may further reduce the interfacial tension.^38^ Alternatively, we evaluated the possibility to initiate the fusion of pH sensitive SUVs to the droplet periphery in a sequential manner, i.e. following the production of SUVs-containing droplets. Towards this end, the preformed SUV-containing droplets were exposed to a surfactant-oil mixture supplemented with 36 mM acetic acid (Figure 2B). A rapid assembly of dsGUVs was observed upon the exposure to the acidic oil conditions. Interestingly, we observed a reduction in size following the introduction of acid, where large droplets failed to support additional mechanical stress associated to the oil substitution.

**Figure 2:**
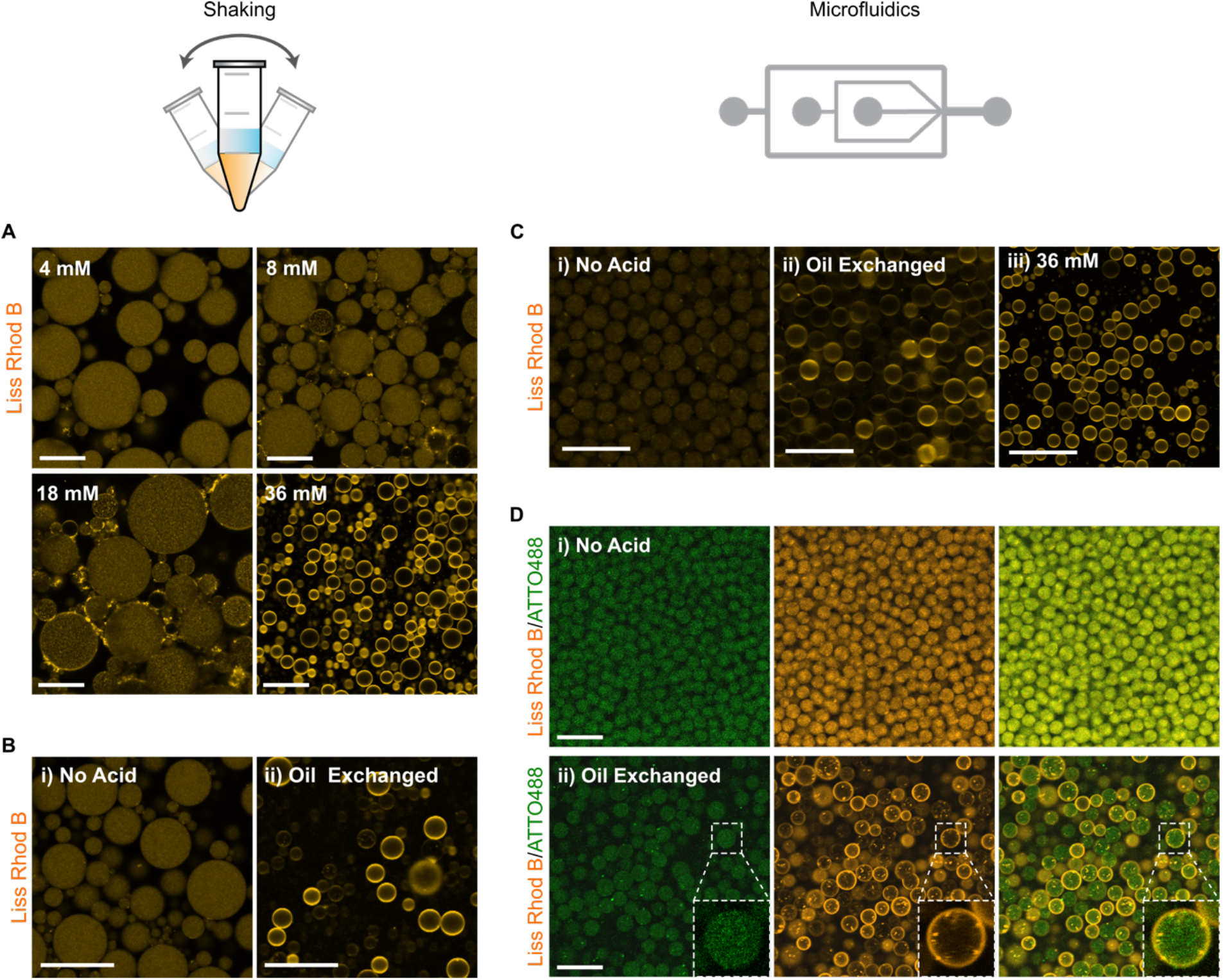
Assembly of dsGUVs by acidification from the oil-surfactant phase via bulk shaking method or microfluidic technology. (**A**) Representative confocal fluorescence images of pH sensitive SUVs (DOBAQ/DOPG/DOPC/Liss Rhod B labelled-DOPE (60/20/19.5/0.5 mol%)) within W/O droplets stabilized by 1.4 wt% PEG-based fluorosurfactant and 10 mM Krytox in HFE-7500 oil that contains various concentration of acetic acid (4, 8, 18 and 36 mM). Droplets were produced by bulk shaking method and the aqueous phase consisted of 1.5 mM SUVs in 50 mM KH_2_PO_4_/K_2_HPO_4_, 75 mM KCl, pH 7.4. Scale bars, 50 μm. (**B**) Following droplets production by a bulk shaking method (**i**), the oil-surfactant mixture was exchanged by the acidic oil-surfactant mix supplemented with 36 mM acetic acid, thus provoking the rapid assembly of a supported lipid bilayer at the droplet periphery (**ii**). Scale bar, 50 μm. (**C**) Sequential assembly of dsGUVs via a microfluidic mechanical splitting module by entrapping SUVs in droplet with an oil-surfactant mix without supplementing acetic acid (**i**) and following the substitution of the oil phase by an acidic oil containing 36 mM acetic acid (**ii**). Alternatively, assembly of dsGUVs produced by microfluidic splitting can be achieved through the direct usage of an acidic oil containing 36 mM acetic acid as continuous phase (**iii**). Scale bars, 25 μm. To minimize droplet coalescence in acidic oil condition, the oil phase contained 3 wt% PEG-based fluorosurfactant, and 10 mM Krytox and 35 mM acetic acid in HFE-7500. (**D**) Assembly of multicompartment dsGUVs by microfluidic mechanical splitting through post-production acidification. Droplets encapsulated two SUV populations: 1.5 mM SUVs composed of DOBAQ/DOPG/DOPC/Liss Rhod B labelled-DOPE (60/20/19.5/0.5 mol%) and 1 mM of Q_pa_DOPE/ATTO 488-labelled DOPE (99.5/0.5 mol%), both in 50 mM KH_2_PO_4_/K_2_HPO_4_, 75 mM KCl, pH 7.4. The oil-surfactant mixture was composed of 3 wt% PEG-based fluorosurfactant, 10 mM Krytox in HFE-7500. (Top). Following the production and collection of the dsGUVs, the oil-phase was substituted by an acidic oil, initiating the rapid and selective fusion of DOBAQ SUVs to the droplet periphery (Bottom). Scale bars, 25 μm.

Following the successful assembly of dsGUVs in bulk conditions by applying a pH-trigger from the oil phase, we evaluated if the use of an acidic oil could empower and facilitate the direct translation of this approach towards small channel geometry microfluidics. We observed that both approaches, either post-production acidification or the direct use of an acidic oil, enable the reliable production of dsGUVs by mechanical splitting (Figure 2C; Supplementary video S2). Interestingly, due to their inherent and homogeneous small size, no significant difference in droplet size was observed before, and after oil substitution, thus reinforcing the idea that larger droplets do not support additional mechanical stress during oil exchange. In addition, the use of an acidic oil still enables the selective recruitment of pH sensitive SUVs to the droplet periphery to allow the assembly of compartmentalized dsGUVs (Figure 2D). By using mechanical splitters, two population of SUVs were co-encapsulated within W/O droplets possessing an average diameter of 7.7 ± 0.9 µm (*n = 212;* Fig 2D Top; FigureS7), and 8.5 ± 1.4 µm (*n = 181;* Fig 2D Bottom; Figure S7) before and after introduction of the acidic oil phase, respectively. Thus, the use of acetic acid - which is soluble in both the water phase and the fluorinated oil phase - can modulate the pH of the W/O droplets to mediate the assembly of dsGUVs encapsulating pH sensitive SUVs.

The pH-mediated assembly of compartmentalized dsGUVs depends on two interconnected parameters: (1) the buffering capacity of the aqueous phase and (2) the pK_a_ of the pH sensitive lipid. Since the use of Krytox, a fluorinated carboxylic acid^39^, will lead to a concomitant acidification of the droplet lumen, we investigated if the use of either low pH citrate buffer or acetic acid in the oil phase could be omitted. Towards this end, we reduced the buffering capacity of the aqueous phase from 50 to 10 mM phosphate buffer and supplemented 140 mM KCl in order to match the osmolarity and buffering capacity of a standard phosphate buffer saline (e.g. 1x PBS). Note, in order to minimize potential interactions in between population of SUVs and the droplet periphery, KCl was favored over NaCl due to the reduced interaction of K^+^ ions with phospholipids^40^. To probe the effect of Krytox on the pH of the droplets, fluorescein was encapsulated within W/O droplets stabilized by 2.5 mM PEG-based fluorosurfactant in HFE-7500 in absence of Krytox. A partitioning assay of the PEG-based fluorosurfactant showed a minimal amount (∼ 46 µM) of Krytox impurity (Figure S8) which was considered negligible. Fluorescein has an intricate sensitivity to pH, its fluorescence intensity decreases in acidic environment (Figure 3A).^41^ The pH of the water phase in presence of fluorescein was varied by adjusting the KH_2_PO_4_/K_2_HPO_4_/K_3_PO_4_ ratio within the range of pH 5 to 8. Droplets were then imaged by CLSM, where the fluorescence intensity showed a linear correlation with the droplet’s inner pH (Figure 3B-C). Then, droplets containing 10 mM KH_2_PO_4_/K_2_HPO_4_ and 140 mM KCl at pH 7.4 were generated by supplementing various concentrations of Krytox (2.5; 5.0; 7.5 and 10 mM) to an oil-surfactant mixture containing 2.5 mM PEG-based fluorosurfactant in HFE-7500. In these conditions, we observed a drastic reduction of the droplet pH, reaching 5.1 at 10 mM Krytox (Figure 3B). Importantly, in such acidic conditions, the DOBAQ containing SUVs typically assemble to generate dsGUVs as previously observed herein when citrate buffer, or acetic acid was employed. We confirmed the successful assembly of dsGUVs by encapsulating pH sensitive SUVs in 10 mM KH_2_PO_4_/K_2_HPO_4_ and 140 mM KCl at pH 7.4 in W/O droplets with solely implementing 10 mM Krytox as acid source (Figure S9).

**Figure 3:**
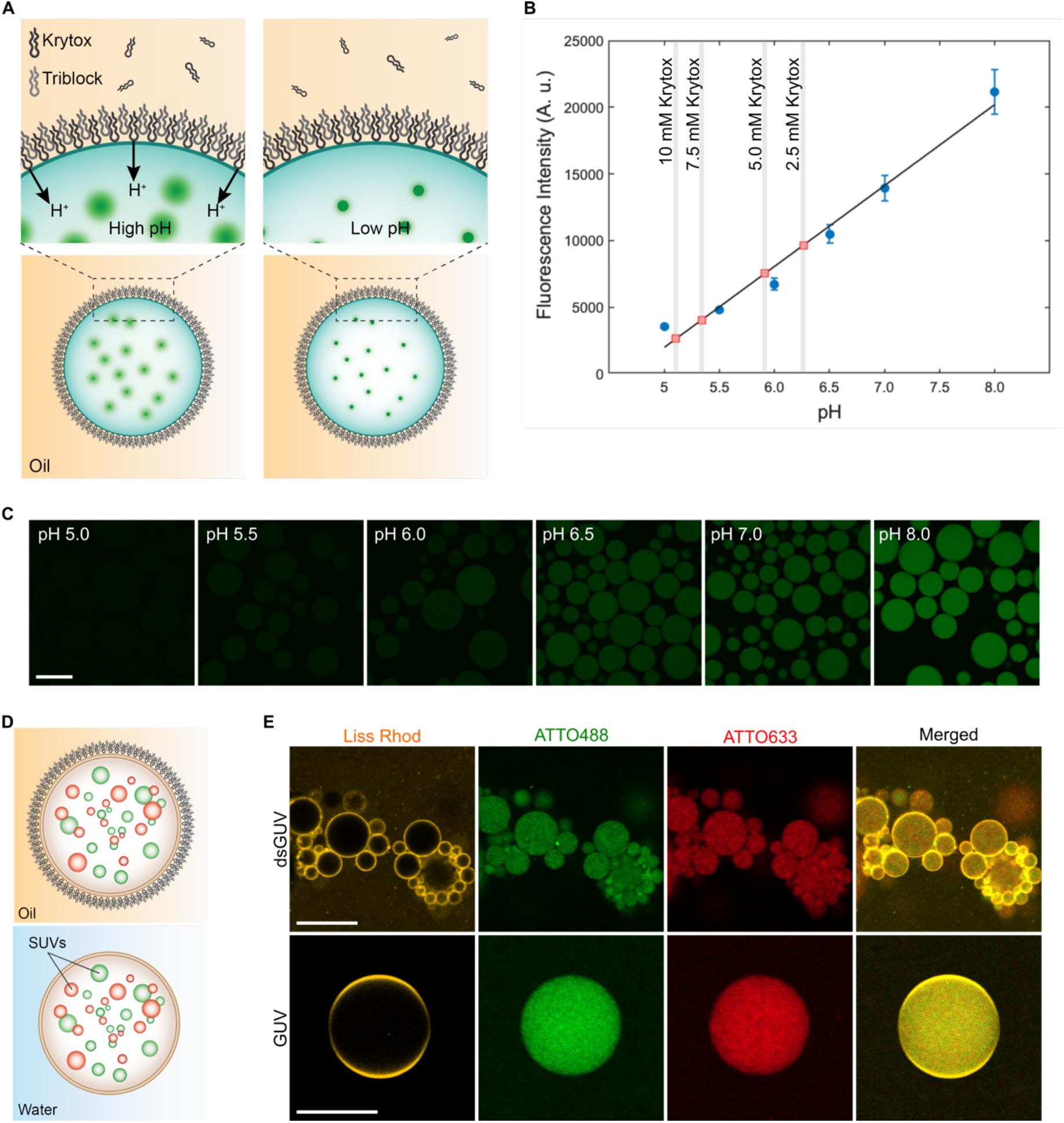
(**A**) Schematic representation of the fluorescein-based detection of concomitant acidification of the W/O aqueous core by the addition of Krytox in the oil-surfactant mixture. At physiological pH (left image), the fluorescence of fluorescein is maximal, while upon acidification the fluorescence is diminished (right image low pH). (**B**) Calibration curve of the mean fluorescence intensity excited at 488 nm of W/O droplets encapsulating 1 μM fluorescein in 10 mM KH_2_PO_4_/K_2_HPO_4_ and 140 mM KCl at various pH. W/O droplets were generated by manual shaking to produce the emulsion, by employing an oil-surfactant mixture composed of 2.5 mM PEG-based florosurfactant and various Krytox concentration (2.5; 5.0; 7.5 and 10 mM, represented by a vertical lines) in HFE-7500. A partitioning assay (see Supplementary information) detected a Krytox contamination of 46 µM into the PEG-based fluorosurfactant, which was considered negligible (Figure S8). The calculated pH of droplet population at each Krytox concentration are depicted by the red squares. Mean ± S.D are presented (*n* = 50 droplets). (**C**) Representative CLSM images of the fluorescein-containing W/O droplets produced at various pH. Scale bar: 100 μm. (**D**) Schematic representation of the generation of multicompartment dsGUVs through the co-encapsulation of different SUVs populations and release in physiological conditions. (**E**) CLSM images presenting the self-assembly of compartmentalized droplet-stabilized GUVs achieved via shaking. (**Top**) dsGUVs generated after the encapsulation three SUV populations: 1.5 mM pH sensitive SUVs composed of DODMA/DOPG/DOPC/DMG-PEG/Liss Rhod B PE (30/15/50.5/4/0.5 mol%), (2) 1 mM of redox sensitive SUVs composed of Q_pa_DOPE/ATTO488-labelled DOPE (99.5/0.5 mol%) and (3) 1mM of negatively charged SUVs composed of DOPG/DOPC/ATTO633-labelled DOPE (30/69.5/0.5 mol%). All SUVs were prepared in 10 mM KH_2_PO_4_/K_2_HPO_4_, 140 mM KCl, pH 7.4. Acidification of the droplet lumen was achieved through the direct presence of Krytox in the oil-surfactant mixture composed of 2.5 mM PEG-based fluorosurfactant, 7.5 mM Krytox in HFE-7500. Scale bar, 25 µm. (**Bottom**) Released GUV in physiological conditions presenting the homogeneous distribution of the inner compartments. Scale bar, 10 µm.

Along with the reduction of the buffering capacity, we produced SUVs incorporating other pH sensitive lipids exhibiting a greater pK_a_ than DOBAQ, which would thus modulate their charge at higher pH. We incorporated DODMA, a synthetic pH sensitive lipid possessing a pK_a_ of 7.6 once incorporated in lipid nanoparticles.^28^ Interestingly, we measured a pK_a_ of 8.2 for DODMA by a TNS assay when incorporated into SUVs, highlighting the impact of the lipid environment on the pK_a_ of the lipid (Figure S10).^42^ By substituting DOBAQ for DODMA, we showed the selective self-assembly of an endomembrane system incorporating different type of compartments with a reduced concentration of Krytox of 7.5 mM when 10 mM KH2PO4/K2HPO4 and 140 mM KCl at pH 7.4 was used as aqueous phase (Figure 3D-E). Herein, functional SUVs, incorporating the redox-sensitive lipid Q_pa_-DOPE labelled with ATTO488, and a passive SUVs labelled with ATTO633 were all encapsulated within dsGUVs by a pH-trigger. Moreover, zwitterionic SUVs (i.e. solely DOPC containing SUVs) were also successfully encapsulated with our pH-mediated approach, thus expanding its potential for the generation of functional multicompartment synthetic cells (Figure S11). These results further highlight the potential of the pH-triggered assembly of dsGUVs to build multi-functional synthetic eukaryotes possessing stimuli-responsive compartments. To summarize, the use of an external source of acid (i.e. from the oil phase) can be employed to either trigger or directly assemble compartmentalized dsGUVs through simple emulsification (i.e. shaking) or microfluidic platforms. Moreover, additional control and flexibility can be introduced into the system while applying a pH trigger to assemble compartmentalized dsGUVs: (1) the buffering capacity and; (2) the pK_a_ of the pH sensitive lipids.

### Usage of pH improve the SUVs to GUVs conversion

As opposed to conventional lipids, which are mostly zwitterionic in nature, pH sensitive lipids grant the capacity to modulate the surface charge of SUVs in a relevant physiological pH window. This charge modulation was used to initiate the assembly of dsGUVs through the reduction of the droplet pH. One of the drawbacks of the charged-mediated assembly of GUVs, which is also related to its advantage, is the use of W/O droplets stabilized by fluorosurfactants. These droplets act as molecular template, where their size dictates the final size of the GUVs while providing important mechanical stability. To release the assembled GUVs into physiological conditions, the droplets must be destabilized, i.e. the stabilizing surfactant must be removed. This destabilization process is achieved through the addition of an excess of small, and poorly stabilizing surfactant such as 1*H*,1*H*,2*H*,2*H*-perfluoro-1-octanol, promoting droplet coalescence and hence, the release of the GUVs into the water phase. To be successful and effectively released from the droplet, molecular interactions between the droplet interface and the lipids must be the lowest in order to minimize mechanical stresses. The typical use of Krytox and Mg^2+^ ions to recruit and fuse SUVs to the droplet periphery corresponds to a strong ionic interaction which cannot be easily altered or dynamically modulated to maximize GUV production. Herein, we envision that the external modulation of the surface charge may facilitate and improve the release efficiency of dsGUVs to isolate free standing GUVs.

Toward this aim, we compared the release efficiency of dsGUVs produced by either the standard procedure using 10 mM Mg^2+^ ions^4, 23, 32^, or assembled by a pH-trigger. For direct comparison, a SUV formulation, composed of DOBAQ/DOPG/DOPC/DOPE/Liss Rhod B-labelled DOPE (30/20/39.5/10/0.5 mol%) was produced. Importantly, this formulation possessed an optimized DOBAQ/DOPG ratio, which enable both the use of pH, or Mg^2+^ ion to mediate the assembly. SUVs were encapsulated into W/O droplet stabilized by 2.5 mM PEG-based fluorosurfactant and 10 mM Krytox in HFE-7500. For the Mg^2+^ ions mediated assembly of dsGUVs, the aqueous phase was supplemented with 10 mM MgCl_2_ and 30 mM Tris buffer pH 7.4. We observed that without Mg^2+^ ions SUVs did not fuse to the periphery, thus confirming no pH-mediated assembly in 30 mM Tris buffer when 10 mM Krytox was used (Figure S12). In the case of pH-mediated assembly, the aqueous phase was supplemented with 10 mM KH_2_PO_4_/K_2_HPO_4_, 140 mM KCl, pH 7.4, and employed the same oil-surfactant mixture. Following their production and incubation, dsGUVs were released with an osmotically matching buffer at pH 7.4. CLSM imaging revealed the striking increase in absolute number of released GUVs when pH was employed compared to Mg^2+^ (Figure 4A, Figure S13). In the case of pH-mediated assembly, we postulated that the interactions between the Krytox and the lipids at the droplet interface were minimized due to the release in pH 7.4. At physiological pH, the surface charge of the dsGUVs becomes more negative as the DOBAQ lipid became a zwitterion at pH > 4.35. This increase in negative charge leads to a reduction of the electrostatic interaction between lipids and Krytox at the droplet periphery, and may even promote electrostatic repulsion. When dsGUVs were released at pH 5, a reduced number of GUVs was observed, which corroborate the presence of strong remaining interactions between Krytox and DOBAQ lipids at pH 5 compared to pH 7.4.

**Figure 4:**
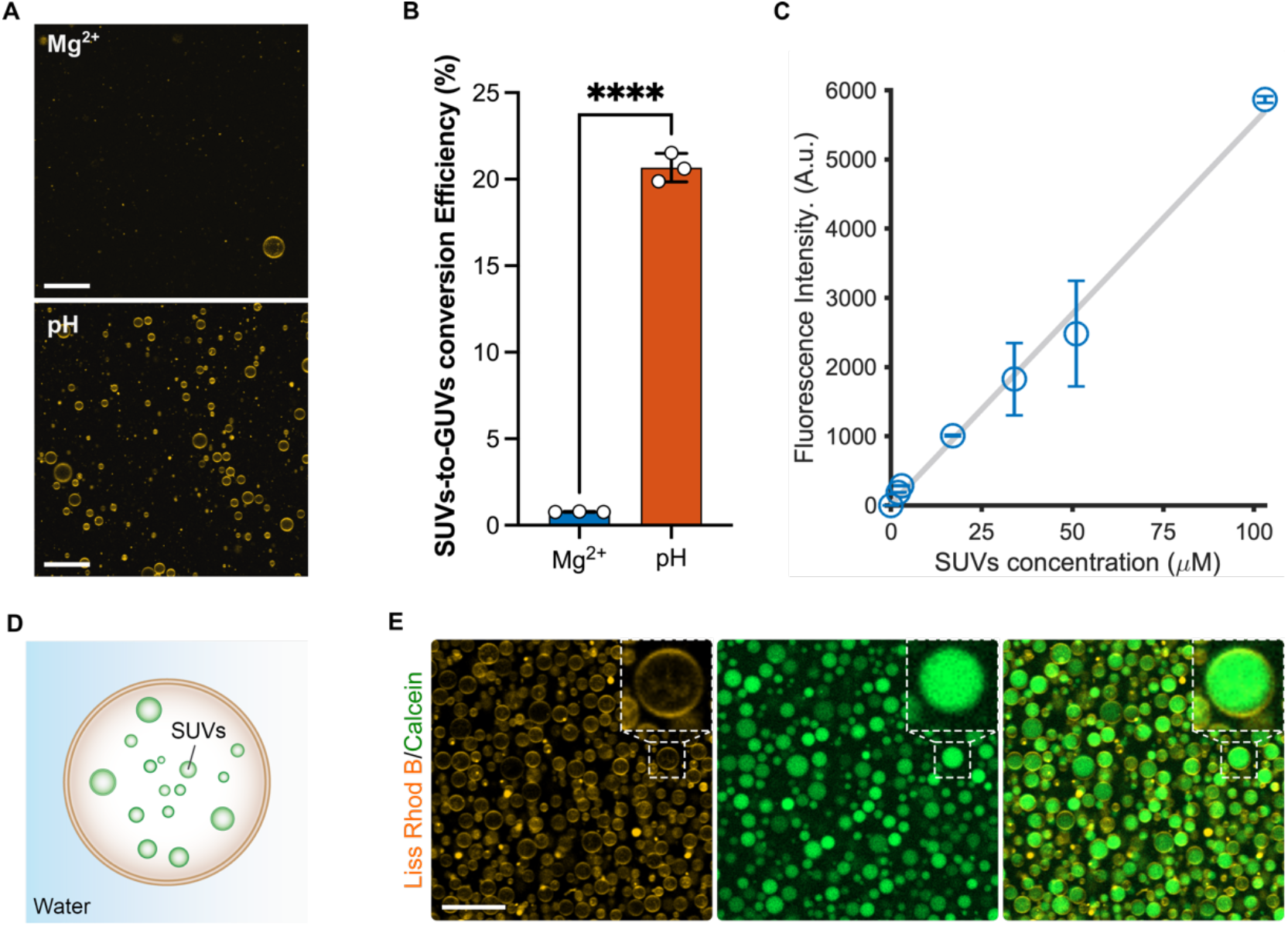
(**A**) Representative CLSM images of randomly selected locations within the observation chamber of released GUVs produced from precursor SUVs containing DOBAQ/DOPG/DOPC/DOPE/Liss Rhod B-labelled DOPE (30/20/39.5/10/0.5 mol%). GUVs were assembled with either the presence of 10 mM MgCl_2_ (referred as Mg^2+^) or at pH 5 (referred as pH) through the acidification of the W/O droplets from the oil phase by Krytox. In the case of Mg^2+^-mediated assembly, the aqueous phase was supplemented with 10 mM MgCl_2_ and 30 mM Tris buffer pH 7.4. In the case of the pH-mediated assembly, the aqueous phase was 10 mM KH_2_PO_4_/K_2_HPO_4_, 140 mM KCl, pH 7.4. W/O droplets were produced by manual shaking to achieve an emulsion. Droplets were stabilized by 2.5 mM of PEG-based fluorosurfactant, and 10 mM Krytox in HFE-7500, with a water to oil ratio of 1:2. Scale bars: 50 μm. (**B**) SUVs-to-GUVs conversion efficiency of GUVs prepared by Mg^2+^- and pH-mediated assembly as evaluated by fluorescence spectroscopy. Mean ± S.D. are presented (*n* = 3). Data were analyzed using an unpaired *t*-test. ****P<0.0001. The SUVs to GUVs conversion efficiency corresponds to *N*_*GUVs*_ */ N*_*GUVs,theoretical*_, where *N*_*GUVs*_ is the total number of GUVs released from the emulsion, and *N*_*GUVs theoretical*_ is the theoretical total number of GUVs possible to produced based on the number of SUVs within the W/O droplets. Lipid concentration of samples was measured by fluorescence according to the signal of Liss Rhod B labelled-DOPE contained in GUVs in a mixture of 1:1 isopropanol:water, with the aid of a calibration curve of the precursor SUVs in isopropanol:water presented in (**C**). Mean ± S.D. are presented (*n* = 3), R^2^ = 0.9912. (**D**) Scheme presenting the formation of free-Standing multicompartment GUVs in physiological conditions. (**E**) Multicompartment GUVs encapsulating 1 mM calcein-loaded Q_pa_DOPE SUVs (100 mol%). The GUVs, were assembled by the pH mediated approach from precursor SUVs composed of DOBAQ/DOPG/DOPC/DOPE/Liss Rhod B-labelled DOPE (30/20/39.5/10/0.5 mol%) employing mechanical splitter microfluidics for W/O droplet production to achieve higher release efficiency, and improve polydispersity. Scale bar: 25 µm.

To further quantify the efficiency of GUVs production between pH and Mg^2+^ ions mediated assembly, we measured what we referred as the SUVs-to-GUVs conversion efficiency (Figure 4B). The SUVs to GUVs conversion efficiency corresponds to the ratio *N*_*GUVs*_ */ N*_*GUVs*_,_*theoretical*_, where *N*_*GUVs*_ is the measured number of GUVs released from the emulsion, and *N*_*GUVs*_,_*theoretical*_ is the theoretical total number of GUVs possible to be produced using the provided lipids during assembly. The number of produced and released GUVs was assessed by fluorescence spectroscopy in a 1:1 isopropanol:water mixture and compared to a calibration curve of the precursor SUVs in 1:1 isopropanol:water (Figure 4C), where the increase in fluorescence was associated to the presence of GUVs rather than lipid aggregates, as shown the CLSM images in figure 4A. Here, the use of an organic solvent to solubilize the lipids into the water phase was essential in order to compare the calibration curve generated from SUVs and GUVs. Results revealed that the SUVs-to-GUVs production efficiency is 20-fold greater when a pH trigger was used over Mg^2+^ ions with an average SUVs-to-GUVs conversion efficiency of 20 %, and also achieved high conversion efficiency with commercially available PEG-based fluorosurfactant (Supplementary note 1; Figure S13). When translated to a microfluidic platform to achieve better polydispersity, we observed a tremendous improvement in production efficiency, thus achieving a high yield generation of compartmentalized GUVs (Figure 4D-E). In summary, the pH-mediated assembly showed improve production efficiency compared to the standard Mg^2+^ ions mediated assembly, while also empowering the generation of multicompartment GUVs.

### pH-mediated assembly of multicompartment GUVs with an actin-cytoskeleton

One of the key aspects of bottom-up synthetic biology relies on the capability to reconstruct cellular functions within cell-sized compartments in order understand fundamental cellular processes with a minimal set of variables and components. Among these essential components, the actin cytoskeleton possesses a pivotal role in a plethora of cellular processes such as adhesion, cellular polarization, migration, division and differentiation.^43^ Its reconstruction into cell-sized vesicles in presence of an endomembrane system is thus an important step in order to reproduce and investigate various cellular functions in synthetic eukaryotes with increased complexity. Up to now, protein-encapsulated dsGUVs in presence of Mg^2+^ ions showed poor production efficiency, and depended on the isoelectric point P_i_ of the protein. Negatively charged proteins, such as actin^44^, may hinder the charge mediated assembly of the lipid bilayer in presence of high Mg^2+^ ions concentration (i.e. 10 mM). To palliate this issue, microfluidic platforms incorporating pico-injectors^45^ can be used to sequentially reconstruct an actin cytoskeleton^4^, but release of actin-containing dsGUVs remained challenging.

Toward this goal, we applied the pH-triggered assembly of GUVs, which allows for improved production efficiency, avoid the needs of high concentration of Mg^2+^ ions and enable the generation of multicompartment GUVs. As a first step, we co-encapsulated pH sensitive SUVs with actin filaments (F-actin) in a W/O emulsion generated by shaking. Upon acidification of the droplets by Krytox, we observed the recruitment and fusion of SUVs at the periphery, while the F-actin remained located within the droplet lumen (Figure S14). The F-actin network was homogeneously distributed within the droplet lumen due to reduced interactions in between the F-actin and Krytox at low concentration of Mg^2+^. Note, a leakage of the Liss Rhod B-labelled DOPE lipids into the fluorinated oil phase under these experimental conditions was observed. The leakage is attributed to the strong stochiometric association of Rhodamine B to Krytox which effects the retention of dyes within W/O droplet as a function of salt concentration, buffer and Krytox concentration.^32, 46, 47^ Following their production, the F-actin containing dsGUVs were successfully released into physiological conditions (Figure 5A-B). We observed that the released GUVs were typically smaller in size compared to the corresponding dsGUVs, suggesting two possible case of figures: First, large vesicles containing F-actin may hardly tolerate the bulk release process, leading to release of intraluminal F-actin into the aqueous buffer visible by CLSM a to bottom of the observation chamber (Figure S15). Secondly, vesicles may shrink due to slight changes in osmotic pressure.

**Figure 5:**
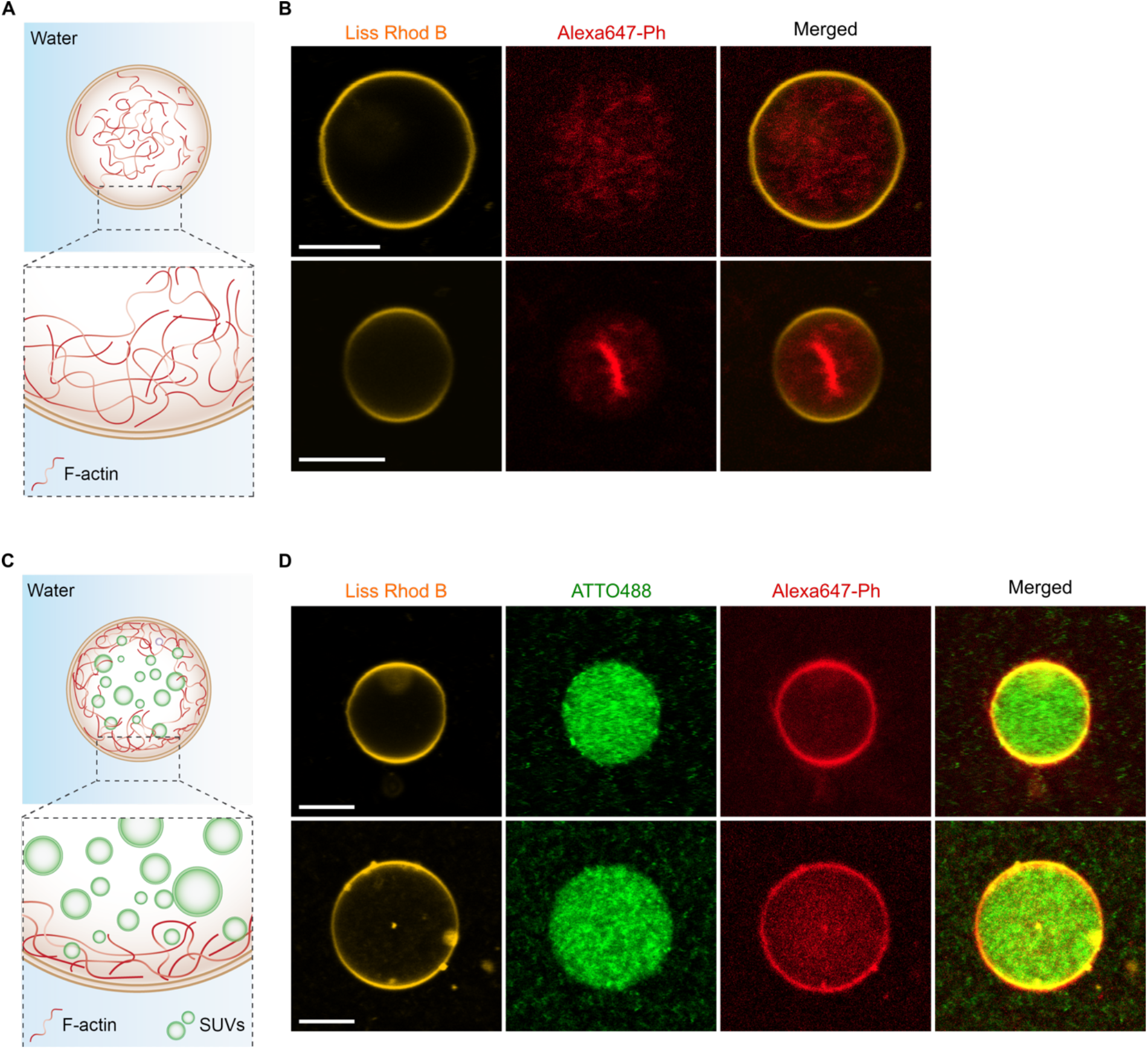
Reconstruction of an F-actin cytoskeleton in absence and in presence of an additional endomembrane system inside a synthetic eukaryote assemble through a pH-trigger within W/O droplet, and release in physiological condition. (**A**) Scheme presenting the encapsulation of F-actin within a free-standing GUV assembled by a pH-trigger presenting the homogeneous distribution of F-actin within the lipid vesicle. (**B**) CLSM images of two representative cases of free-standing GUVs depicting either the relatively homogeneous distribution of the F-actin filaments, or the confinement of the actin bundles at the center of the vesicles. Prior their release, the GUVs were assembled by co-encapsulating pH sensitive SUVs containing DOBAQ/DODMA/DOPG/DOPE/DOPC/DMG-PEG/Liss Rhod B-labelled DOPE (15/15/15/10/40.5/4/0.5 mol%), and encapsulate 5 µM of F-actin labelled Alexa647-Phalloidin in surfactant-stabilized W/O droplet produced by vortexing. Scale bars: 10 µm. (**C**) Scheme presenting the encapsulation of both F-actin and SUVs imitating an endomembrane system within a free-standing GUV assembled by a pH-trigger. (**D**) Representative CLSM images of two free-standing GUVs depicting a homogeneous distribution of the negatively charged inner compartment within the vesicle, whereas F-actin preferentially accumulate at the periphery of the lipid vesicle. The GUVs were assembled by supplementing 1 mM of negatively charged SUVs composed of DOPG/DOPC/ATTO488-labelled DOPE (30/69.5/0.5 mol%) to the mixture presented in (**B**), and encapsulated within surfactant-stabilized W/O droplets produced by vortexing. Scale bars: 10 µm.

Our results show that the produced GUVs typically retain their homogeneous distribution of F-actin (Supplementary Video S3), or in few cases, exhibited a further accumulation of F-actin toward the center of the vesicle as previously reported by Weiss and coworkers.^4^ This interesting difference and improvement in assembly can be first explained by the production method (pH-triggered, choice of PEG-based fluorosurfactant and reduced Mg^2+^), but also by the introduction of additional electrostatic interactions resulting from changes in pH. Actin monomers possess an isoelectric point at pH ≈ 5.4^43^, meaning that upon acidification by Krytox, actin will possess a slight excess of positive charges. In this case, both the SUVs and the F-actin would possess an excess of positive charges, thus minimizing their interaction within each other. Moreover, the pH sensitive SUVs are expected to diffuse more rapidly to the negatively charged droplet interface owed to their smaller size. By assuming a 3D Brownian motion of 100 nm SUVs encapsulated within a 15 µm diameter W/O droplet, and by applying the Stoke-Einstein law of diffusion for a spherical particle, an SUV located at the center of the droplet would reach the droplet interface in ≈ 400 ms, corresponding to a diffusion coefficient *D*_*SUV*_ of 24 µm^2^/s (Supplementary Note 2). F-actin, on the other hand, possesses a translational diffusion coefficient *D*_*actin*_ typically inferior to 1 µm^2^/s when located within two thin walls.^48^ Even though *D*_*actin*_ is expected to be higher in unbound fluids (i.e. within the W/O droplet), the diffusion of F-actin is greatly affected by its length, which in our case, will significantly be reduced by the presence of long micrometer-scaled filaments. Consequently, SUVs will diffuse faster than the F-actin at pH ≈ 5 toward the droplet interface to promote the assembly of dsGUVs.

Interestingly, when compartments were co-encapsulated with F-actin and pH sensitive SUVs in W/O droplets, we observed accumulation of F-actin at the vesicle’s periphery, even in absence of any (bio)chemical linkers^49^. In addition, the compartments showed a homogeneous distribution within the droplet-stabilized vesicle (Figure S16), and after their release in physiological conditions (Figure 5C-D). This behavior is associated to the depletion effect, where F-actin organized itself at the vesicle periphery in presence of compartments. In this spatial configuration, the entropy of the SUVs is maximized, thus corresponding to the most thermodynamically favor structure of the system. Similar observations were previously reported for large particles, which spontaneously adsorbed at the vesicle periphery when co-encapsulated with smaller negatively charged particles.^50, 51^

### Summary and outlook

In summary, the usage of pH as internal, or external trigger to activate the charge-mediated assembly of dsGUVs allows the reconstruction of an endomembrane system in either a bulk assembly or by microfluidics. To achieve a successful formation of GUVs, the method relies on two important criteria: (1) slightly negatively charged SUVs can be efficiently entrapped and (2) the use of a pH sensitive lipid. Moreover, the assembly is dictated by the apparent pK_a_ of the pH sensitive SUVs, which could then be readily optimized through chemical synthesis of desired lipids, where the pK_a_ of synthetic lipids were extensively investigated and tuned in the past decade.^26, 52^ By introducing a pH sensitive motif, the surface charge of the GUVs can be modulated, which is the origin of a fundamental improvement in total number of GUVs produced from the droplet-stabilized approach. Besides improving the production efficiency, the use of pH has empowered the reconstruction of an F-actin cytoskeleton without, with an endomembrane system. This corresponds to a very basic advancement in bottom-up synthetic biology employing solely lipid-based vesicles. Interestingly, we observed a drastic change in the behavior of F-actin in presence of compartments due to the depletion effect, highlighting the possibility to observe and investigate emergent properties resulting from the combination of different cellular module. Still, further experiments and optimization would be required to investigate the behavior of F-actin in presence of molecular crowder, such as SUVs, inside GUVs in greater details. Additionally, adapting the methodology to encapsulate actin monomers, rather than filamentous actin, could offer greater possibility. Perhaps, compartments could be engineer to facilitate or impede actin polymerization and ultimately, recreate cellular motility inside synthetic eukaryotes. The method presented herein could thus catalyze the assembly of more specialized synthetic eukaryotes for biotechnological and medical applications. By entrapping various stimuli-responsive compartments, minimal cross-reactivity and control over the release sequence could be embedded within a single synthetic eukaryote. The assembly of such an endomembrane system within a micron-scale carrier could be engineered to release therapeutics through the vascular walls^53^, while enabling improved functionality, reduced passive leakage, cross-reactivity and flexibility to name a few.

## Author Information

## Acknowledgments

The authors acknowledge funding from the Federal Ministry of Education and Research of Germany, Grant Agreement no. 13XP5073A, PolyAntiBak and the MaxSynBio Consortium, which is jointly funded by the German Federal Ministry of Education and Research and the Max Planck Society. They also acknowledge the support from the SFB 1129 of the German Research Foundation and the Volkswagenstiftung (priority call ‘Life?’). F.L. acknowledges the support of the Alexander von Humboldt Foundation. F.L. would like to thank Dr. Dimitris Missirlis for fruitful discussion regarding GUVs production and quantification. F.L. would also thank Tobias Abele for its help regarding the Fiji script for droplet localization. K.J. thanks the Carl Zeiss Foundation for financial support. J.P.S. acknowledge funding from the Deutsche Forschungsgemeinschaft (DFG, German Research Foundation) under Germany’s Excellence Strategy via the Excellence Cluster 3D Matter Made to Order (EXC-2082/1 – 390761711) and the Gottfried Wilhelm Leibniz Award. The Max Planck Society is appreciated for its general support.

## TOC

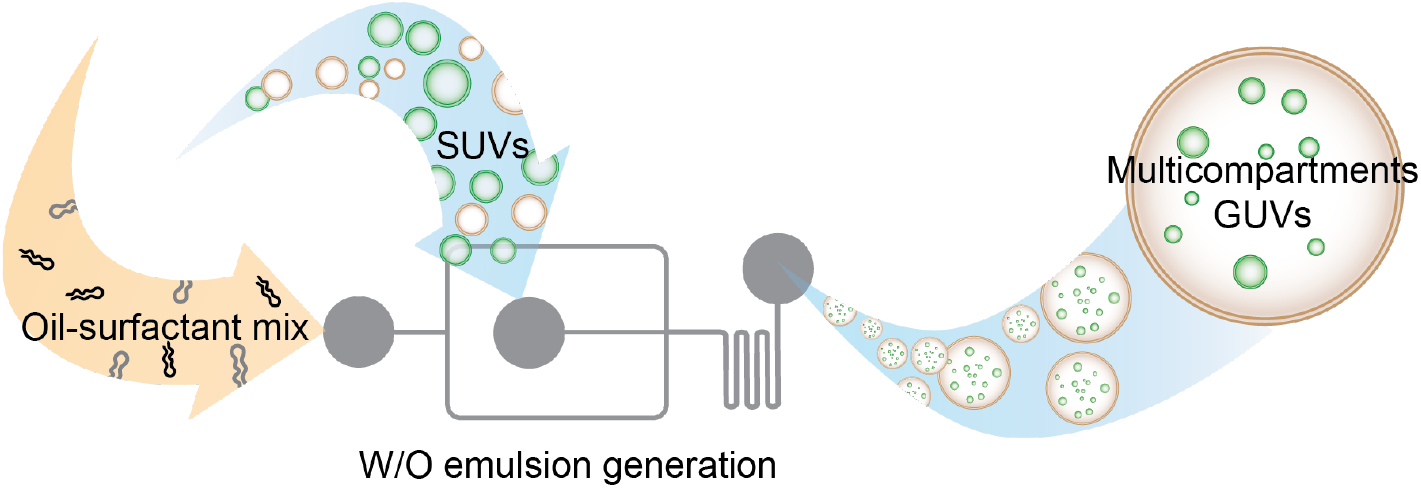

