## Supplementary Information for "pH-Triggered Assembly of Endomembrane Multicompartments in Synthetic Cells"

**Materials and methods:**

ATTO488-labelled 1,2-dioleoyl-sn-glycero-3-phosphoethanolamine (ATTO488-labelled DOPE) was purchased from ATTO TEC (Germany). 1,2-dieoleyloxy-3-dimethylaminopropane (DODMA) was synthetized as described elsewhere.[1] Quinone proprionic acid linked-1,2-dioleoyl-sn-glycero-3-phosphoethanolamine ($Q_{pa}$DOPE) was prepared as reported elsewhere.[2] All other lipids were bought from Avanti Polar Lipids ® as solution in chloroform, and used without further purification. All lipids were stored at -20 °C until needed. All chemicals were bought from Sigma-Aldrich (Germany) and were used without further purification. Bovine serum albumin (BSA) coated glass slides (25x60 $mm^2$ and 18x18 $mm^2$) were prepared by drop coating a solution of 1 mg/mL BSA in 1x DPBS for 15 minutes, dried and rinsed with MilliQ water. The BSA-coated glass slides were directly used to assemble an observation chamber through the usage of double-sided sticky tape.

**General procedure for preparation of small unilamellar vesicles (SUVs)**: SUVs were prepared by lipid film hydration, followed by extrusion. Lipids from stock solutions in chloroform were mixed to the desired molar ratio in a glass vial. Chloroform was evaporated by blowing with a gentle stream of nitrogen to obtain a thin lipid film. To remove residual traces of organic solvent, the vial was desiccated under vacuum for a period of two hours. The dry lipid film was then rehydrated through the addition of a solution of 10 mM $KH_2PO_4/K_2HPO_4$, 140 mM KCl, pH 7.4 (if not mentioned otherwise), at a final lipid concentration of 6 mM. The lipid film was swelled for 20 minutes, and vortexed 30 seconds to trigger the rapid formation of multilamellar liposomes. The lipid suspension was extruded through a 100 nm polycarbonate track-etch membrane (Whatman), 21 times at a temperature at least 5 °C above the $T_m$ of the lipids, 25 °C in most of our case, with a miniextruder (Avanti Polar Lipids, Inc.). The resulting SUVs were stored at 4 °C until needed for up to 3 days or used immediately for generating dsGUVs.

**Calcein loaded-$Q_{pa}$DOPE SUVs**: 5 mg of $Q_{pa}$DOPE was dissolved in 5 mL of chloroform in a round bottom flask and evaporated under vacuum with a rotary evaporator for one hour. The dried lipid film was re-hydrated with a solution of 50 mM calcein dissolved in 50 mM $KH_2PO_4/K_2HPO_4$, 75 mM KCl, pH 7.4 to a final lipid concentration of 1 mg/mL. The lipid film was aged for one hour with occasional vortexing every 15 minutes, followed by five cycles of freeze-thawing in a dry ice/acetone bath. The lipid suspension was extruded through a 100 nm polycarbonate track-etch membrane (Whatman), 21 times at room temperature with a miniextruder (Avanti Polar Lipids, Inc.). Following extrusion, un-encapsulated calcein was removed by spin column filtration. Briefly, Sephadex-G50 resin fine (GE healthcare Bioscience) was swelled for at least three hours in 50 mM $KH_2PO_4/K_2HPO_4$, 75 mM KCl, pH 7.4. The resin was added over a glass-wooled plugged 2 mL syringe until the resin completely filled the syringe, and then compacted by centrifugation (2 minutes, 1000g). 200 uL of liposomal solution was added to the column, and centrifugated for 10 minutes at 50 g. Eluted calcein-loaded liposomes were then expelled and collected from the column by centrifuging 2 minutes at 1000g. The resulting unilamellar calcein-loaded SUVs were stored at 4 °C until needed for up to 7 days. Note that in the case of calcein loaded SUVs, extrusion was performed through a 100 nm membrane to facilitate and improve the purification step achieved by size exclusion.

**Synthesis of triblock-co-polymer fluorosurfactant:** The synthesis of fluorosrufactant, composed of PFPE-$PEG_{1500\ MW}$-PFPE was adapted from Scanga and coworkers.[3]

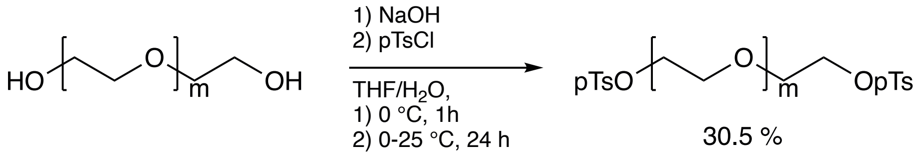


***$PEG_{1500}$ ditosylate* (1)*:*** Sodium hydroxide (13.32 g, 333 mmol, 4.0 eq.) was dissolved in water (103 mL) under ice cooling. Tetrahydrofuran (200 mL) was added to the solution and the mixture was allowed to cool down to 0 °C. PEG 1500 (125 g, 83.3 mmol, 1.0 eq.) was added in small portions, so that the temperature did not rise above 5 °C. Afterwards, the mixture was allowed to warm up to room temperature and stirred for one hour. After the reaction mixture was cooled to 0 °C again, *p*-toluenesulfonic acid (36.4 g, 191 mmol, 2.3 eq.) dissolved in tetrahydrofuran (220 mL) was added dropwise and care was taken that the temperature did not rise above 5 °C. The reaction mixture was stirred for 18 hours, allowing the temperature to rise to room temperature. The organic layer was separated and the solvent was removed under reduced pressure. After dissolving the crude product in ethyl acetate (750 mL), the solution was washed with 90 vol% brine (100 mL). The solution was dried over magnesium sulfate and filtered. Afterwards, the solvent was removed under reduced pressure. The tosylated product **1** was received as a white solid (46.0 g, 25.4 mmol, 30.5 %). $^1$H NMR (400 MHz, $CDCl_3$): δ = 2.385 (s, 6H, H6), 3.578 (m, H3, H2), 4.090 (t, J = 4.4 Hz, 4H, H1), 7.283 (d, J = 4.0 Hz, 4H, H5), 7.729 (d, J = 4.0 Hz, 4H, H4).

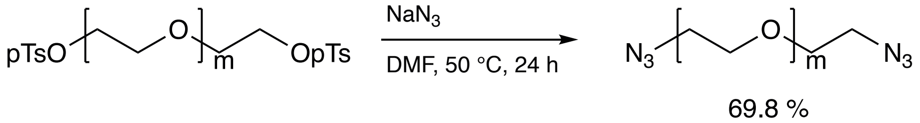


***$PEG_{1500}$ diazide* (2)**: Compound **1** (20.0 g, 11.05 mmol, 1.0 eq.) was dissolved in dimethylformamide (30 mL) and sodium azide (1.58 g, 24.3 mmol, 2.2 eq.) was added. The mixture was stirred first for 90 minutes at room temperature followed by 18 hours at 50 °C. As the reaction progresses, the mixture becomes more and more turbid. The mixture was filtered, and the solvent was removed by co-evaporation with toluene under reduced pressure. Afterwards, the crude product was resuspended in ethyl acetate (200 mL), filtered and washed with 90 vol% brine (50 mL). The obtained solution was dried over magnesium sulfate, then filtered and the solvent was removed under reduced pressure. The diazide substituted product **2** was received as a white solid (11.9 g, 7.71 mmol, 69.8 %). $^1$H NMR (400 MHz, $CDCl_3$): δ = 3.379 (t, J = 5.2 Hz, 4H, H1), 3.634 (m, H2, H3).

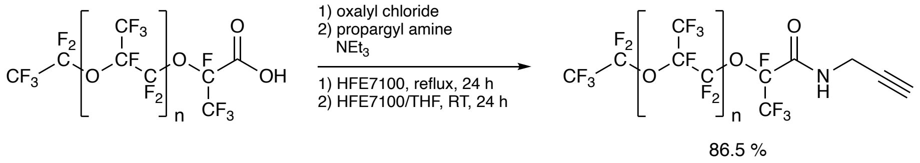

***Propargyl PFPE$_{7000}$*** **(3)**: In the first step, PFPE acid (Krytox FSH, 90.23 g, 12.89 mmol, 1.0 eq.) was filled into a flame-dried flask, degassed under reduced pressure and dissolved in HFE 7100 (150 mL). Oxalyl chloride (3.4 mL, 39.4 mmol, 3.05 eq.) was added and the reaction mixture was refluxed for 18 hours, during which the reaction mixture became cloudy. The solvent and excess of oxalyl chloride were removed under reduced pressure, collecting the removed material in a liquid nitrogen cooling trap. In a second step, the intermediate product was dissolved in HFE 7100 (90 mL). The reaction vessel was equipped with dropping funnel under nitrogen counterflow. The funnel was loaded with propargyl amine (870 µL, 13.5 mmol, 1.05 eq.), triethylamine (2.7 mL, 19.3 mmol, 1.5 eq.) and tetrahydrofuran (35 mL). The propargyl amine solution was added dropwise and the reaction mixture was stirred for 18 hours. The solvent and other volatile reagents were removed under reduced pressure. The obtained crude product was dissolved in HFE 7100 and filtered. After removing the solvent under reduced pressure, the propargyl derivative **3** was received as an orange oil (78.5 g, 11.15 mmol, 86.5 %). As NMR solvent, a mixture of $C_6F_6$:$C_6D_6$ 88:12 was used. $^1$H NMR (400 MHz, $C_6D_6$): δ = 2.186 (t, J = 2.8 Hz, 1H, H3), 4.135 (t, J = 2.8 Hz, 2H, H1), 6.602 (s, 1H, NH).

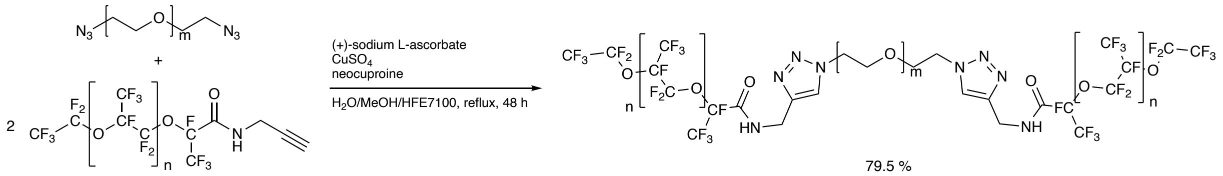


**PFPE$_{7000}$-PEG$_{1500}$-PFPE$_{7000}$ triazole linked triblock fluorosurfactant (SynFS):** Compound **3** (2.31 g, 1.49 mmol, 1.05 eq.), sodium ascorbate (112 mg, 568 µmol, 40 mol%), copper(II) sulfate pentahydrate (70.9 mg, 284 µmol, 20 mol%) and neocuproine (94.8 mg, 455 µmol, 32 mol%) were dissolved in water (15 mL) and methanol (15 mL). Propargyl PFPE 7000 (20.0 g, 2.84 mmol, 2.0 eq.) was dissolved in HFE 7100 (30 mL) and added to the aqueous solution. The reaction mixture was refluxed for 48 hours. The reaction mixture was cooled to room temperature. Afterwards, it was transferred in a separation funnel and carefully overlaid with methanol (15 mL). The two-phase system was swiveled without mixing. The methanol layer became orange and was removed after saturation. This procedure was repeated until the methanol phase did not become orange anymore. If the volume of fluorinated oil phase was decreased as much that the swiveling was not efficient anymore, HFE 7100 was added. Afterwards, the fluorinated phase was dried over magnesium sulfate. The crude product was first filtered through Celite and then through a 0.45 µm PFTE syringe filter. The solvent was removed under reduced pressure. The triblock copolymer surfactant was received as a highly viscous substance (17.6 g, 1.13 mmol, 79.5 %). $^1$H NMR (400 MHz, $C_6D_6$): δ = 3.637 (m, H3), 3.970 (s, 4H, H2), 4.588 (s, 4H, H1), 4.678 (m, 4H, H6), 7.976 (s, 2H, H4), 8.433 (s, 2H, NH). The synthetized fluorosurfactant will be referred as SynFS.

**Microfluidic Device Fabrication:** Microfluidic devices were fabricated using polydimethylsiloxane (PDMS) and soft lithographic procedure. All microfluidic devices were designed using the computer-aided design (CAD) software QCAD-pro (RIbbonSoft, Switzerland). Briefly, a thin layer of negative photoresist SU8-3005 (MicroChem, USA) was spin coated (Laurell Technologies Corp., USA) at 1000 rpm on a 2 in. silicon wafer to produce a uniform 10 μm think layer. Then, wafers were soft-baked on a hot plate at 65 °C for 1 minute, then ramped and held at 95 °C for 3 minutes. The CAD designs were directly exposed to the photoresist through a Tabletop Micro Pattern Generator μPG 101 (Heidelberg Instruments, Germany) employing the writing mode II. The exposure conditions were set to 70 mW for the output laser power, and 25% for the pixel pulse duration. Following the exposure, the wafers were baked on a hot plate at 65 °C for 1 minute, then ramped and held at 95 °C for 5 minutes. Next, mr-DEV 600 (MicroChemicals, Germany) was used to remove the unexposed SU8 resists from the wafer, and hard baked in an oven at 150 °C for 15 minutes. After baking, the wafer was placed in a Petri dish and served as a mold for downstream PDMS fabrication. PDMS (Sylgard 184, Dow Corning, USA) was well mixed with the curing agent at a 10:1 ratio, degassed, poured onto the wafer, and cured for 2 hours at 65 °C in an oven. The PDMS was then cut, and carefully peeled off from the mold. Holes, serving as inlets and outlets were generated using a 0.5 mm biopsy punch (Harris Uni-Core, Ted Pella, Inc.). Following punching, holes in the PDMS were cleaned with isopropanol and pressurized nitrogen gas to remove residual PDMS particles. Glass slides (24 x 60 $mm^2$, Carl Roth, Germany) were sequentially cleaned using heptane and isopropanol, and thoroughly dried with pressurized nitrogen gas. To bound the PDMS to a coverslip, the device and a clean glass slide were activated using an oxygen plasma (TePla, Germany; 0.4 mbar, 200 W, 35 seconds exposure) and brought in contact with gentle pressure. To strengthen the attachment between the PDMS and the glass slide, devices were heated 1 hour at 65 °C in an oven. Sigmacote (Sigma-Aldrich, Germany), was passed through the channels to render their surface hydrophobic.

**General pH mediated assembly of multicompartment GUV in bulk**: Initially, we prepared an aqueous solution containing two population of SUVs; 1.5 mM of pH-sensitive SUVs composed of DOBAQ/DOPG/DOPE/DOPC/DMG-PEG/Liss Rhod B DOPE (30/20/6/39.5/4/0.5 mol%) in 10 mM $KH_2PO_4/K_2HPO_4$, 140 mM KCl, pH 7.4, and 1 mM of the SUVs to be entrapped, typically composed of DOPG/DOPC/ATTO488 labelled-DOPE (30/69.5/0.5 mol%) in 10 mM $KH_2PO_4/K_2HPO_4$, 140 mM KCl, pH 7.4. Then, an oil-surfactant mix composed of 2.5 mM of SynFS and 10 mM PFPE-carboxylic acid (Krytox-157FSH, MW 7000-7500 g/mol, DuPont, Germany) in HFE-7500 (3M, Germany) was prepared, and filtered through a 0.22 μm polycarbonate filter. As an alternative, the SynFS may be substituted by commercially available perfluoropolyether-polyethylene glycol (PFPE-PEG) fluorosurfactant (Ran Biotechnologies, Inc.) at a final concentration of 1.4 wt/wt%. However, we observed a substantial decrease in release efficiency when pH was employed as strategy of self-assembly as opposed to $Mg^{2+}$. Then, 50 μL of the SUVs containing aqueous solution was layered on top of 100 μL of the oil-surfactant mix, and vortexed vigorously for 30 seconds. The visible and persistent milky-like emulsion located above the excess of oil phase indicates the formation of stable water-in-oil droplets. In these conditions, the pH sensitive SUVs were preferentially recruited at the droplet periphery and fused to from a spherical supported lipid bilayer resulting in the entrapment of the other SUVs population within the formed droplet-stabilized GUV. To release the GUVs from the oil-surfactant phase, 75 μL of 1x DPBS (2.7 mM KCl, 1.47 mM $KH_2PO_4$, 8.1 mM $Na_2HPO_4$, 138 mM NaCl, Gibco, ThermoFisher Scientific), or an osmolarity-matched buffer was slowly added on top of the droplet emulsion. In order to destabilize the droplets, 75 μL of 1*H*,1*H*,2*H*,2*H*-perfluoro-1-octanol (PFO) acting as a destabilizing agent was slowly added on top of the aqueous phase. The sample was gently rolled, and left at 4 °C overnight. After incubation, the milky emulsion disappeared and led to a transparent aqueous layer on top of the oil-surfactant mixture. The top aqueous layer, containing the released GUVs, was carefully collected with a micropipette, while avoiding the collection of oil, and either directly transferred into a BSA-coated observation chamber for direct fluorescence imaging, or stored in a new test tube at 4 °C until needed for up to 3 days. Although direct release is possible, we observed that the release efficiency (i.e. the absolute number of GUVs per mL) was significantly improved by storing the dsGUVs at 4 °C for a period of at least 2h hours, and ideally overnight, prior the addition of the release buffer and the destabilizing agent. As a general rule, the pH mediated assembly was optimized to a water: oil ratio of 1:2, while we noted a maximal release efficiency when a water:release buffer (and PFO) of 1:1.5 ratio was used. The assembly can easily be scaled up and down, based on the experimental needs, without significant loss in release efficiency as long as these ratios are kept constant.

**pH mediated assembly of multicompartment GUV by microfluidics:** Stable water-in-oil droplets were generated at the flow-focusing T-junction by employing the oil-surfactant mixture and the aqueous phase described above as continuous and disperse phase, respectively. Fluids were introduced within the microfluidics via PTFE tubing (0.3 mm I.D., 0.6 mm O.D., BOLA Tubing) with a Pressure Controller (OB1 Mk3; Elveflow). Pressure was adjusted with the ESI software (v3.01.13; Elveflow,). Typical pressure for flow-focusing T-junction were between 500 – 750 mbar for both the disperse and continuous phase, while a pressure of 2 bar for both disperse and continuous phase was needed for the mechanical splitters. The droplets production was assessed trough an axio table-top inverted microscope (Zeiss, Germany) via a 10x LD-A-Plan objective (NA = 0.25) equipped with a high-speed camera EoSens CL (Mikrotron GmbH). Following the production of the droplets, the resulting dsGUVs were incubated for a period of at least 2h at 4 °C, or ideally overnight, prior their release. The release procedure remains the same as for the bulk production described above. In general, the volume of droplet produced was in the range of 100 μL.

**Actin preparation:** Actin was purified from acetone powder from New Zealand white rabbit skeletal muscle, based on the method of Pardee and collegues[4], and modified according to Kron and coworkers[5], and stored in 2 mM Tris-HCl, 0.2 mM $CaCl_2$, 0.2 mM ATP, 0.005 % wt/v $NaN_3$, and 0.2 mM DTT at pH 8, at −80 °C. Actin monomers were labeled with Phalloidin-Alexa647 (Sigma-Aldrich) by mixing 72 μL actin monomers with 10 μL of 10x Actin Polymerisation Buffer (APB; 20 mM Tris-HCl, pH 8.0, 500 mM KCl, 20 mM $MgCl_2$, 10 mM NaATP) and 18 μL of 2x Actin Buffer (AB; 50 mM Imidazole, pH 7.4, 50 mM KCl, 2 mM EGTA, 8 mM $MgCl_2$). The actin monomers were left at room temperature to polymerize for 30 minutes. Subsequently, 10 μL (20 μL in MeOH, evaporated to ~ 10 μL) of Phalloidin-Alexa647 (10 units) were added to the solution. The resulting phalloidin-Alexa647-labelled F-actin was stored at -80 °C until needed.

**F-actin encapsulation with and without inner compartment into GUVs:** The reconstruction of an F-actin cytoskeleton within GUVs in presence of absence of inner compartment was achieved by co-encapsulation pH sensitive SUVs and polymerized F-actin into W/O droplets stabilized by a PEG-based fluorosurfactant in presence of low amount of $Mg^{2+}$ ions. First, pH sensitive SUVs containing DOBAQ/DODMA/DOPC/DOPG/DOPE/DMG-PEG/Liss Rhod B-labelled DOPE (15/15/44.5/15/6/4/0.5 mol%) were produced in by lipid film hydration with actin buffer (AB; 25 mM imidazole, 25 KCl, 1 mM EGTA, 4 mM $MgCl_2$, pH 7.4) at a concentration of 6 mM of lipid, and extruded as described in the general procedure above. Inner compartments composed of DOPC/DOPG/ATTO488-labelled DOPE (79.5/20/0.5 mol%) were also prepared by lipid film hydration in AB, at a final concentration of 6 mM of lipid, and extruded. To assemble the synthetic eukaryote possessing an F-actin cytoskeleton, pH sensitive SUVs were gently mixed with F-actin labelled with Phalloidin-Alexa647 in AB at a final concentration of 1.5 mM and 5 µM respectively in a 1.5 mL Eppendorf. Alternatively, 1 mM of inner compartments may be supplemented into the aqueous phase in order to recreate an endomembrane system. Then, 50 µL of the SUVs F-actin mixture was layered onto 100 µL of a fluourous phase composed of 2.5 mM PEG-based fluorosurfactant, and 2.5 mM Krytox in HFE-7500 and rapidly emulsified by vortexing to generated droplet-stabilized GUVs. The resulting dsGUVs were stored at 4 $^{o}C$ for 2 hours prior their release. To release the dsGUVs into physiological condition, 75 µL of AB was added to the Eppendorf, followed by 75 µL of PFO, and left undisturbed at 4 $^{o}C$ until complete disappearing of the milky emulsion is achieved. Free-standing GUVs were then imaged by CLSM in a sealed BSA-coated observation chamber.

**Confocal fluorescence microscopy**: Confocal fluorescence imaging was performed on a Zeiss LSM 800 confocal microscope (Carl Zeiss AG, Germany) with a 20X (NA = 0.8) Plan-Apochromat air objective (Carl Zeiss AG, Germany). The pinhole aperture was set to one Airy Unit, and experiments were performed at room temperature. A digital offset of 2500 (16 Bits images) was added to each channel to facilitate thresholding. Collected images were brightness and contrast adjusted, and analyzed with Fiji.[6]

**FRAP Measurements**: The FRAP measurements were perform on a Zeiss LSM800 confocal microscope equipped with a 63X (NA = 1.4) Plan-Apochromat oil objective (Carl Zeiss AG, Germany). The dsGUVs were sealed within a BSA-coated observation chamber, and place in a thermostatic chamber at 25 $^{o}C$. Two circular areas of 2.5 µm radius were defined as probed area at the bottom of each dsGUVs determinate by z-stack profiling beforehand: (1) as bleaching spot, and (2) as reference spot (unbleached) for data correction. Using the bleaching, experimental regions, and time series options in Zeiss Zen software (Zen v2.3), 10 images (laser power, 1.0 %) were recorded prior bleaching (100 iteration; laser intensity, 100%), and 100 images after bleaching (laser intensity, 1.0 %) as depicted in Figure 1B. A 561 nm laser (excitation of Liss Rhod B) was used for the FRAP measurements, with a pinhole aperture set to one Airy Unit. The 247 x 247 pixels images were recorded with an integration of 71.85 ms per image. The diffusion coefficient was extracted from the acquired images using an adapted MatLab (MathWorks, Inc.) code as described previously[7], where a non-linear least-square fit was applied to the normalized fluorescence intensity from the recovery phase. Details and codes are presented in the Supplementary note 2.

**Zeta-Potential measurements**: The zeta potentials of SUVs and free-standing GUVs were diluted to 50 µM phosphate buffer saline at the desired pH. Measurements were performed on a Malvern ZetaSizer Nano ZS in a folded capillary zeta cell (Malvern). The refractive index of the dispersant was set to 1.330, and the viscosity to 0.882 cP with a dielectric constant of 79. The $\kappa \cdot \alpha$ value was set to 1.5. The refractive index on the colloids was set to 1.42, which matched the refractive index of the GUVs. A 5 minutes equilibration time was used prior every measurement for thermal stabilization. For each experimental condition, samples were measured in triplicate, with a minimal of 10 runs per measurements. The maximal voltage was set to 25 V to minimize potential oxidation/reduction effect of the lipid with the capillary electrodes.

**FRET assay**: Lipid mixing between the pH sensitive/positively charged liposomes and negatively charged liposomes acting as compartment was investigated by fluorescence resonant energy transfer (FRET) between donor and acceptor dyes as a function of pH. The FRET probe NBD-PE and Liss Rhod B PE, acting as donor and acceptor, respectively, were formulated within the same pH sensitive/cationic liposomes resulting in a quenching of the NBD signal due to FRET toward Rhod B PE lipids located in the close vicinity. Upon lipid mixing with the negatively charged liposomes, the mean distance between the NBD and Rhod B lipids increases, resulting in an a rapid unquenching of the NBD fluorescence. The pH sensitive SUVs were composed of DOBAQ/DOPG/DOPC/NBD-PE/Liss Rhod B PE (60/20/18/1/1 mol%), cationic liposomes were composed of DOTAP/DOPC/NBD-PE/Liss Rhod B PE (30/68/1/1 mol%) while anionic liposomes were formulated with DOPC/DOPE/DOPG/Cholesterol (45/20/20/15 mol%). All liposomes were prepared at a final lipid concentration of 10 mM by thin-film rehydration. Lipids films were rehydrated with 10 mM Tris-HCl, 50 mM NaCl at pH 8.5, swelled 20 minutes, vortexed 30 seconds and extruded 21 times 100 nm etch-polycarbonate membrane (Whatman) at room temperature with a mini extruder (Avanti Polar Lipids). To investigate the lipid mixing at various pH, different buffers were prepared to cover a pH range from pH 3.0 to pH 8.5, with 0.5 pH unit increments and adjusted with HCl/NaOH. All buffers included 50 mM NaCl, and 10 mM of the buffering compound. For pH 3.0 to 5.5, we used sodium acetate; pH 6.0 to 7.0, we used 2-(*N*-morpholino)ethanesulfonic acid (MES), and pH 7.5 to 8.5, we used Tris-HCl. For lipid mixing experiments, 2 µL of donor liposomes (either pH sensitive, or cationic liposomes) were added to 1998 µL of the corresponding buffer, and the baseline fluorescence was recorded ($F_{min}$). Then, 495 µL of the resulting donor liposome solution was mixed with 5 µL of the negatively charged liposomes, mixed, and measured after 10 minutes of incubation at room temperature ($F$). Finally, 180 µL of the resulting solution was mixed with 20 µL of a 1 wt/wt % Triton-X100, gently mixed and measured after 10 minutes ($F_{max}$). The percentage (%) of lipid mixing was calculated by 100 x $(F - F_{min}) / (F_{max} - F_{min})$. Measurement were performed in triplicates with a Tecan Spark plate reader in Flat 96 wells plate OptiPlate Black (Perkin Elmer), Ex/Em = 465/520 nm, number of flashes = 50, Settle time = 150 ms.

**pKa evaluation by 2-(*p*-toluidino)-6-naphtalenesulfonic acid (TNS) assay**: To evaluate the pKa of the different pH sensitive lipids, a TNS- based assay was employed. Briefly, liposomes were formulated with pH sensitive lipids/DOPC (30/70 mol%) in PBS pH 6.5 at a final lipid concentration of 6 mM. A 1 mM TNS solution, in 9:1 (v/v) EtOH:MilliQ water was prepared. Then, liposomes were diluted to 30 µM in presence of a final TNS concentration of 10 µM in 200 µL per well of a 96-well plate with buffers containing 10 mM 4-(2-hydroxyethyl)-1-piperazineethanesulfonic acid (HEPES), 10 mM 4-morpholineethanesulfonic acid (MES), 10 mM ammonium acetate and 130 mM NaCl, whom pH was adjusted in the range of 2.5 to 11 by 0.5 increments with HCl/NaOH. Afterward, the fluorescence of TNS was measured in triplicate at each pH using a Tecan Spark plate reader in a flat 96 wells plate OptiPlate Black (Perkin Elmer), Ex/Em = 321/431 nm, number of flashes = 30, Settle time = 150 ms. Fluorescence of each liposome formulation at the various pH was then corrected, and normalized according to the value at pH 2.5. A sigmoid function was applied with MatLab (MathWorks) to the fluorescence data, and the pKa of the pH sensitive lipid was approximated as the pH at the point of half maximal fluorescence intensity.

**Evaluation of the SUVs-to-GUVs conversion efficiency**: pH sensitive SUVs employed to assemble, and produce free-standing GUVs through a pH-trigger were diluted to 50 µL in buffer, and mixed with 50 µL isopropanol and vortexed to generate standards (2,5, 15, 50, and 100 µM standards final concentration) as presented in Fig. 4A. Then, 10 µL of Free-standing GUVs (released in a total aqueous phase of 100 µL final) were diluted to 50 µL in buffer, and mixed with 50 µL isopropanol and vortexed. Fluorescence intensity of the Liss Rhod B-labelled DOPE lipids was assessed using a Tecan Spark plate reader in a flat 96 wells plate OptiPlate Black (Perkin Elmer), Ex/Em = 535/595 nm, number of flashes = 50, Settle time = 150 ms, volume per well = 100 µL. Samples and standards were measured in triplicate. The SUVs-to-GUVs conversion efficiency (%) was evaluated by the following equation:

$$\%Conversion = 100\% \cdot \frac{N_{GUVs\,Exp}}{N_{GUVs\,Theo}} = 100\% \cdot f \cdot \frac{c_{GUV} \cdot V_{Well}}{c_{Lip} \cdot V_{prod}}$$

Where *f* is a dilution factor, $c_{GUV}$ is the lipid concentration of the free-standing GUVs measured by the calibration curve generated with the precursor SUVs, $V_{well}$ is the well's volume used to assess the fluorescence intensity by a plate reader measurement and $V_{Prod}$ is the total volume of aqueous phase used to generate the W/O emulsion. Further details on calculation and generated of the presented equation are provided in the Supplementary note 3.

**Partitioning assay**: The Krytox contamination in fluorosurfactant was assessed by a partitioning assay. In a first steps, 100 µL Krytox standards in HFE-7500 (0-1 mM Krytox, 0.1 mM steps) and fluorosurfactant samples (1.4 wt/wt% for commercially available PEG-based fluorosurfactant (CFS), or 2.5 mM for the self-synthetized PEG-based fluorosurfactant (SynFS)) in HFE-7500 were prepared. Then, 100 µL of 1 mM Rhodamine 6G solution in LC-MS grade water was layered onto the fluorinated phase. Careful pipetting ensures that no emulsion was generated throught the process. The mixtures of the two immiscible liquid were incubated for at least 48 h, protected from light. Afterwards, 10 µL of the HFE-7500 phases were collected and diluted to 100 µL with HFE-7500. The partition was calculated by measuring the change in absorbance at 530 nm with a Tecan Spark plate reader in a flat 96 wells plate (TPP Techno Plastic Product) as a function of Krytox concentration, as depicted in Fig. 4A. The calibration curve was employed to evaluate the Krytox impurity. The molar percentage (mol%) of Krytox impurity was defined as the ratio of Krytox concentration divided by the correct concentration of PEG-based fluorosurfactants, which excluded the mass of Krytox impurity.

**Data analysis**: Numerical data were analyzed and plotted with various self-written codes in MatLab (2019a, MathWork). Data fitting was performed in with the curve fitting tool box (Mathwork). In all cases, a robust Bisquare regression method was applied. Fluorescence images were analyzed with Fiji. For the pH measurement inside W/O droplets, droplet detection, localization and assessment of their fluorescence intensity was achieved through a self-written macro in Fiji. Statistical analysis was performed with Prism 9 (GraphPad Software). An unpaired *t*-test analysis, while assuming a Gaussian distribution presenting similar standard deviation (parametric without correction) was used to determine the statistical significance of the pH and $Mg^{2+}$-mediated method of assembly (Fig. 4). Alternatively, a two-way analysis of variance (ANOVA), with a Sidak's multiple comparisons test was used to determine the statistical significance of the pH- and $Mg^{2+}$-mediated method of assembly as a function of the PEG-based fluorosurfactant (Fig. S12). The sample size for figures requiring statistical analysis is stated within the corresponding figure caption.

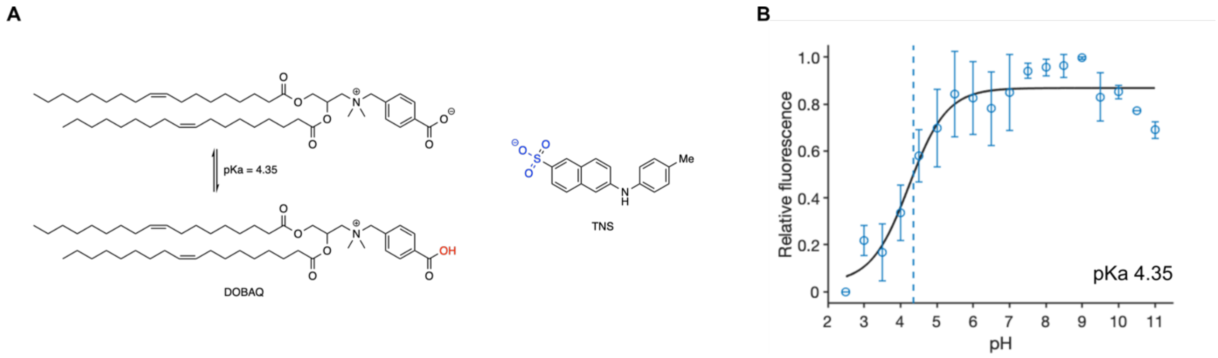


**Figure S1**: Evaluation of the $pK_a$ of DOBAQ by TNS assay. (**A**) Molecular structure of DOBAQ and TNS. The carboxylic group in acidic pH, acting as a hydrogen bond donor is highlighted in red, while the sulfonate group of TNS, acting as a hydrogen bond acceptor is highlighted in blue. (**B**) TNS-based assessment of the pKa value of the pH sensitive SUVs. The pKa is defined as the half-maximum, corresponding to 50 percent of protonated DOBAQ lipid, extracted from a fitted sigmoid curve. The pKa is highlighted by the hatched vertical line. Mean ± S.D. are presented, *n = 3*.

Surprisingly, the TNS response showed a decrease in fluorescence intensity in acidic environment, even though an increase intensity was expected based on previously published experiments.[1, 8-10] The molecular structure of the pH sensitive moieties was shown by Walsh and coworkers to impact its interaction with TNS, mitigating, to some extent, the expected rise in fluorescence from TNS.[11] Compared to most potent pH sensitive lipids, typically possessing tertiary amines[8, 12], DOBAQ possesses a permanent positively charged quaternary amine sterically hindered by a carboxylic acid function located at the forefront of the lipid headgroup (Fig. S1A). As a result, DOBAQ possesses a zwitterionic state in physiological condition, while acidic environment leads to the protonation of the carboxylic acid group rendering DOBAQ positively charged (Fig S1A). We hypothesize that the decrease in fluorescence is associated to the improved capability of DOBAQ to act as an hydrogen bond donor toward the negatively charged TNS at low pH, leading to an apparent increase of the polarity of the local lipophilic environment thus explaining the shape the of TNS assay (Fig S1B).

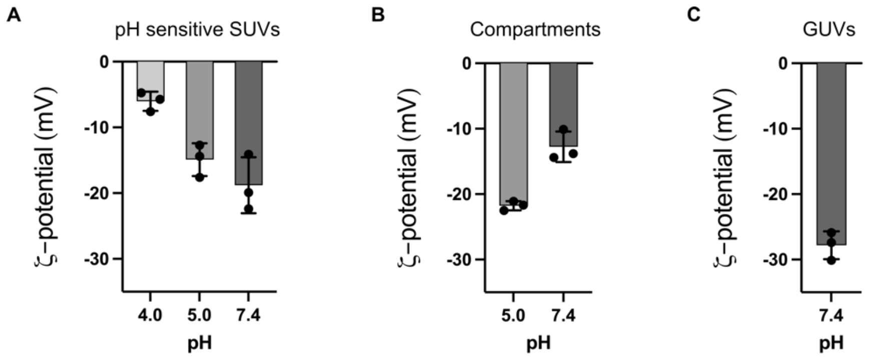


**Figure S2**: Evaluation of the charges of the two population of SUVs as a function of pH, and the resulting free-standing GUVs in physiological condition assembled by a pH trigger. (**A**) the pH sensitive SUVs containing DOBAQ/EggPC/EggPG/Liss Rhod B-labelled DOPE (60/19.5/20/0.5 mol%) and (**B**) the inner compartments SUVs containing EggPC/EggPG/ATTO488-labelled DOPE (79.5/20/0.5 mol%) by zeta-potential measurements. (**C**) Zeta-potential of the free-standing GUVs in physiological condition containing lipids originated from the pH sensitive SUVs in (**A**). In all cases, fluorescently-labelled lipids (Liss Rhod B, and ATTO488) were kept within the formulation to considered their potential effect of the charge of the colloids. Moreover, no interference with the instrument laser (He-Ne, 633 nm) is expected from to the presence of the fluorescent dyes whom possess an excitation wavelength of 561, and 488 nm for the Liss Rhod B and ATTO488, respectively. Mean ± S.D. ($n$ = 3) are presented.

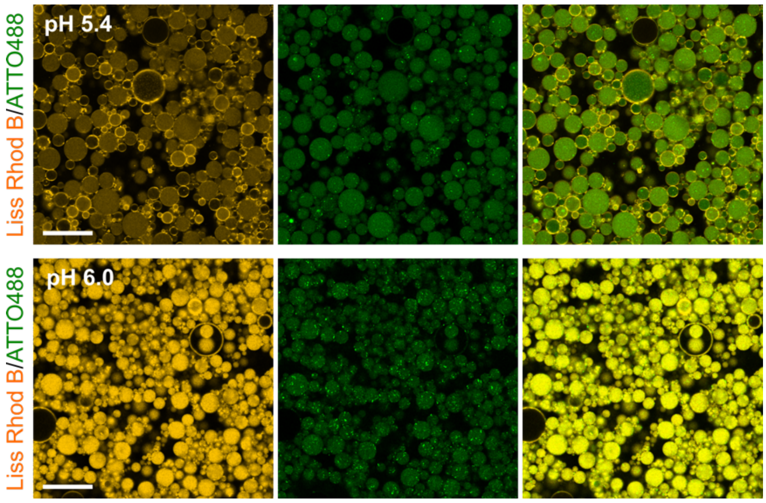


**Figure S3**: Assembly of droplet-stabilized GUVs via shaking in presence of 50 mM citrate buffer at various pH. By decreasing the intra luminal pH with a citrate buffer, pH sensitive SUVs exhibited a preferential fusion to the droplet periphery to generate a supported lipid bilayer. Scale bar, 50 μm. Oil phase: 2.5 mM PEG-based fluorosurfactant, 10 mM Krytox in HFE-7500. The pH sensitive SUVs composed of DOBAQ/eggPG/eggPC/Liss Rhod B-labelled DOPE (60/20/19.5/0.5 mol%) were co-encapsulated with negatively charged SUVs composed of DOPC/DOPG/ATTO488-labelled DOPE (79.5/20/0.5 mol%) within the W/O droplets stabilized by 1.4 wt/wt% fluorosrufactant and 10 mM Krytox. Various 50 mM citrate buffers we used to adjust the pH of the water phase. Scale bars: 50 μm. The acidification of the aqueous phase was achieved through the rapid addition of 100 mM citrate buffer at the desired pH to the aqueous phase right before emulsification. The final citrate concentration in the aqueous phase was 50 mM in all cases.

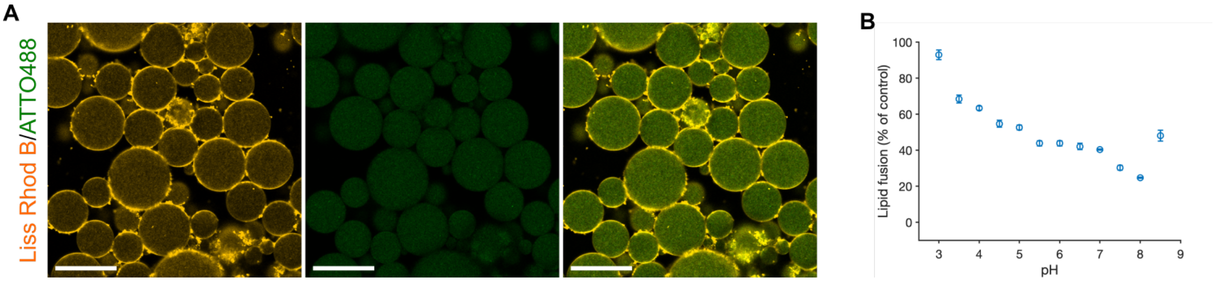


**Figure S4**: Assembly of multicompartment dsGUVs by mixing positively, and negatively charged SUVs population within water-in-oil droplets. (**A**) CLSM images of droplet-stabilized GUVs generated by encapsulating positively charged SUVs composed of DOTAP/DOPC/Liss Rhod B-labelled DOPE (30/69.5/0.5 mol%) and negatively charged SUVs, composed of DOPG/DOPC/ATTO488-labelled DOPE (30/69.5/0.5 mol%). Scale bars; 50 μm. (**B**) FRET assay measuring lipid mixing of positively charged SUVs with negatively charged SUVs. SUVs composed of DOTAP/DOPC/Liss Rhod B-labelled DOPE/DOPE-NBD (30/68/1/1 mol were mixed with unlabeled negatively vesicles at various pH. A relatively constant lipid mixing may be appreciated through, independent of the pH of the environment, which may account for the homogenous fluorescence signal within the dsGUVs lumen presented in (A). Mean ± S.D. are presented, *n = 3.*

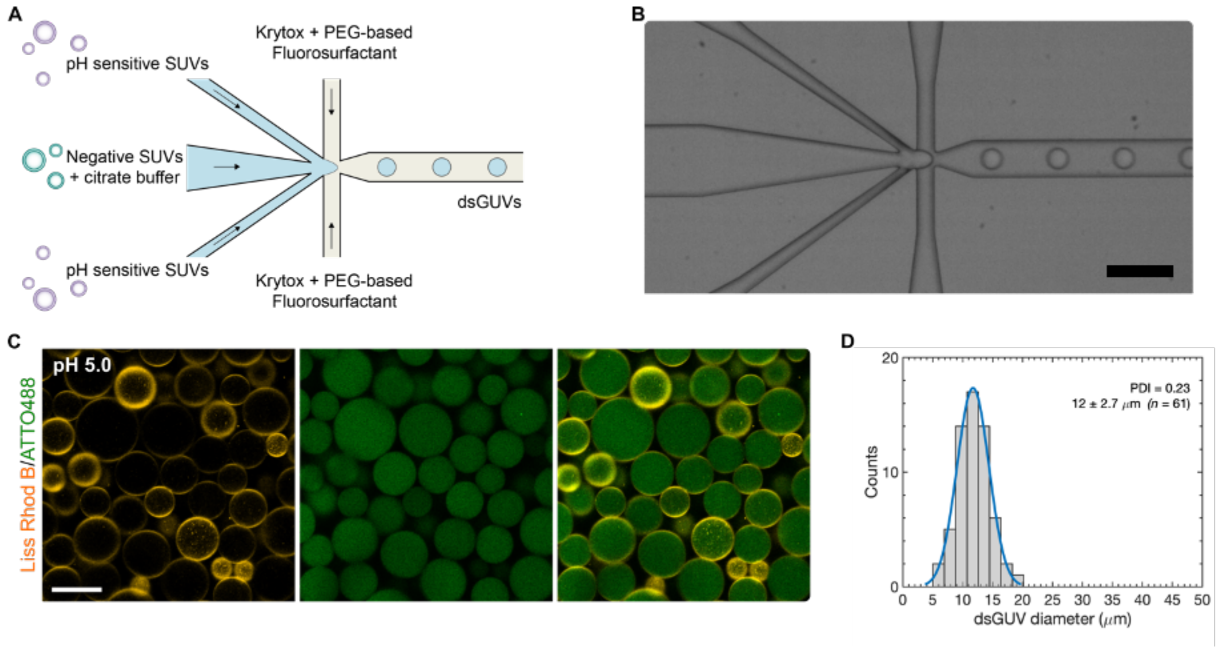


**Figure S5**: Generation of an endomembrane system within droplet-stabilized giant-unilamellar vesicles (dsGUVs) by microfluidics employing two aqueous inlets. (**A**) Schematic representation presenting the production of multicompartment dsGUVs trough the co-encapsulation of a pH sensitive SUVs containing DOBAQ lipids, and an inner compartment SUVs possessing a net negative charge in presence of a citrate buffer at pH 5. The droplets are stabilized through the use of a PEG-based fluorosurfactant and Krytox supplemented to the fluorinated oil-phase. (**B**) Optical micrograph of the microfluidic chip at the flow-focusing junction. Scale bar, 100 µm. (**C**) CLSM images of dsGUVs produced by microfluidics in presence of citrate buffer pH 5. The pH sensitive SUVs, composed of DOBAQ/EggPC/EggPG/Liss Rhod B-labelled DOPE (60/20/19,5/0,5 mol%) showed a preferential accumulation and fusion at the droplet periphery compared to the inner compartment composed of EggPC/EggPG/ATTO488-labelled DOPE (69,5/30/0,5 mol%). Scale bar, 50 µm. The two inlets device was applied to introduce independently: (1) 2 mM of the inner compartment with 100 mM citrate buffer pH 5 and (2) 3 mM of the pH sensitive SUVs. Due to the concomitant dilution of the two aqueous phase prior droplet encapsulation, the concentration was doubled compared to the bulk production achieved by emulsification by shaking. (**D**) Corresponding histogram of the size distribution of the droplets presented in (**C**). The blue lines correspond to the fitted Gaussian distribution. The size distribution was obtained with ImageJ using manual thresholding, and manual measurements of the droplets/dsGUVs diameters. Data were analyzed with MatLab in order extract the Gaussian distribution, and evaluate the polydispersity (PDI) of the produced dsGUVs. During the production, we noted a significant variation of the pressure in between the two inlets, which led to an increase polydispersity in droplet size. We further observed a potential clogging of the inlet channels, resulting from the rapid drop in pH promoting the aggregation of the pH sensitive SUVs.

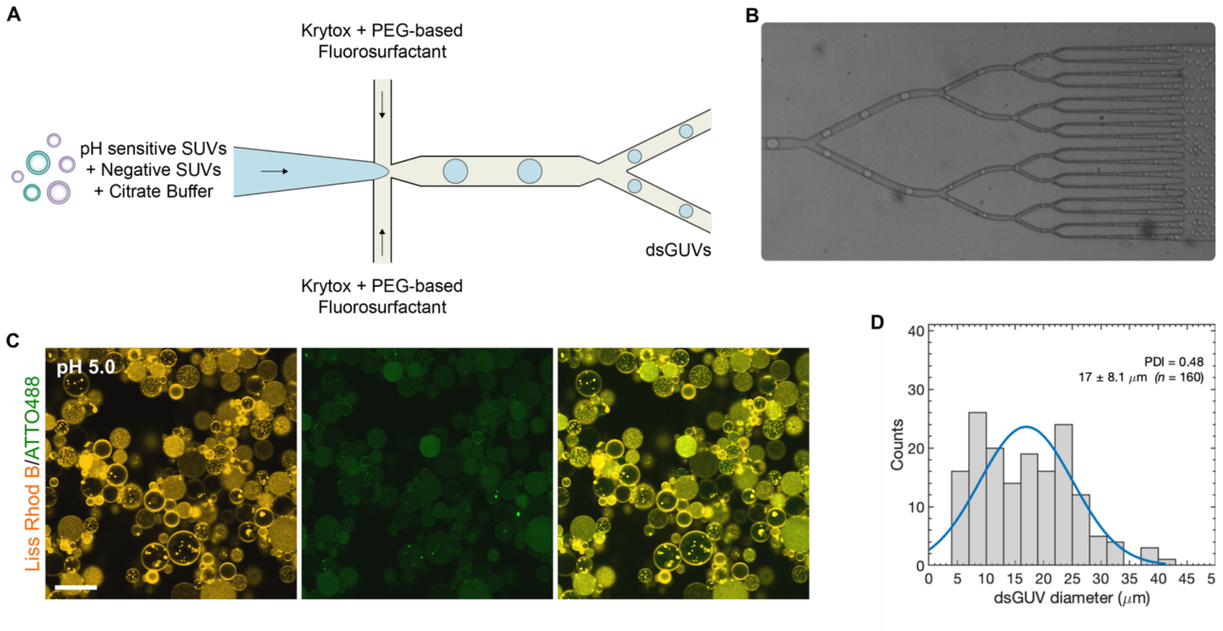


**Figure S6**: Attempt to generate an endomembrane system within small droplet-stabilized giant-unilamellar vesicles (dsGUVs) by microfluidics employing mechanical splitters. (**A**) Scheme presenting the production of multicompartment dsGUVs trough the co-encapsulation of a pH sensitive SUVs containing DOBAQ lipids, and a inner compartment SUVs possessing a net negative charge in presence of a citrate buffer at pH 5. The droplets are stabilized through the use of a PEG-based fluorosurfactant and Krytox supplemented to the fluorinated oil-phase. (**B**) Optical micrograph of the microfluidic chip at the sequential mechanical splitting units exhibiting a concomitant reduction in channel width. (**C**) CLSM images of dsGUVs produced by mechanical splitter in presence of citrate buffer pH 5. The pH sensitive SUVs, composed of DOBAQ/EggPC/EggPG/Liss Rhod B-labelled DOPE (60/20/19,5/0,5 mol%) showed no preferential recruitment and fusion at the droplet periphery compared to the inner compartments composed of EggPC/EggPG/ATTO488-labelled DOPE (69,5/30/0,5 mol%), while also presenting numerous aggregated SUVs. Scale bar, 50 µm. The two inlets, not schematized in (**A**) due to the presence of a 500 µm length channel post inlets but prior the flow-focusing junction, enabled the independent introduction of: (1) 2 mM of the inner compartment with 100 mM citrate buffer pH 5 and (2) 3 mM of the pH sensitive SUVs. Due to the concomitant dilution of the two aqueous phase prior droplet encapsulation, the concentration was doubled compared to the bulk production achieved by emulsification by shaking. (**D**) Corresponding histogram of the size distribution of the droplets presented in (**C**). The blue lines correspond to the fitted Gaussian distribution. The size distribution was obtained with ImageJ using manual thresholding, and manual measurements of the droplets/dsGUVs diameters. Data were analyzed with MatLab in order extract the Gaussian distribution, and evaluate the polydispersity (PDI) of the produced dsGUVs. During the production, we noted a significant variation of the pressure in between the two inlets, which led to an increase polydispersity in droplet size. We further observed a potential clogging of the inlet channels, resulting from the rapid drop in pH promoting the aggregation of the pH sensitive SUVs. Data showed the large polydispersity associated to the aggregation of the SUVs at low pH and the concomitant clogging of the microfluidics channels.

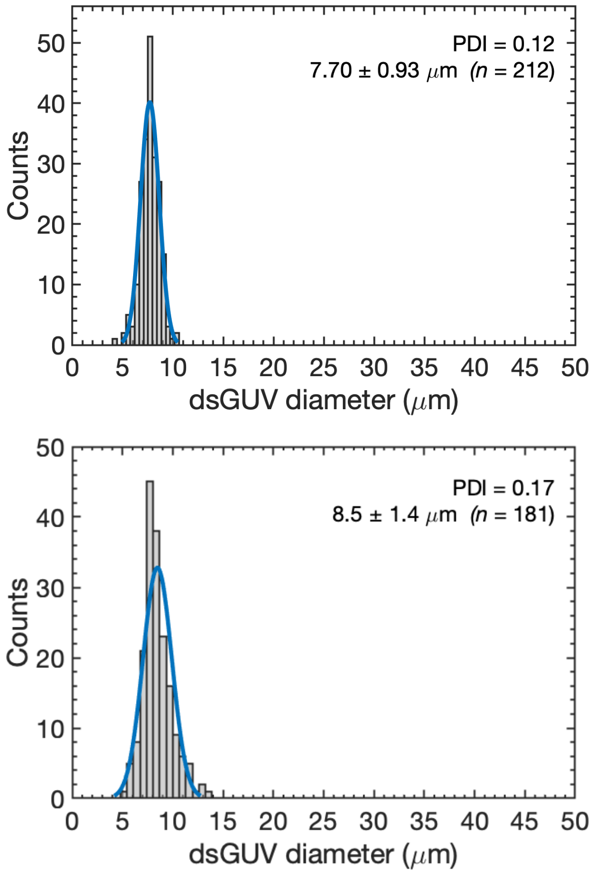


**Figure S7**: Histogram of the size distribution of the droplets (Top), and droplets-stabilized giant unilamellar vesicles possessing an endomembrane system (Bottom) measured by CLSM of the image presented in Fig.2D produced by microfluidic mechanical splitters. Herein, droplets encapsulating various SUVs were produced in absence of supplemented acid, while the dsGUVs assembly was triggered upon exposing the droplets to an acid-containing oil-surfactant phase. The blue lines correspond to the fitted Gaussian distribution. The size distribution was obtained with ImageJ using manual thresholding, and manual measurements of the droplets/dsGUVs diameters. Data were analyzed with MatLab in order extract the Gaussian distribution, and evaluate the polydispersity (PDI) of the dsGUVs.

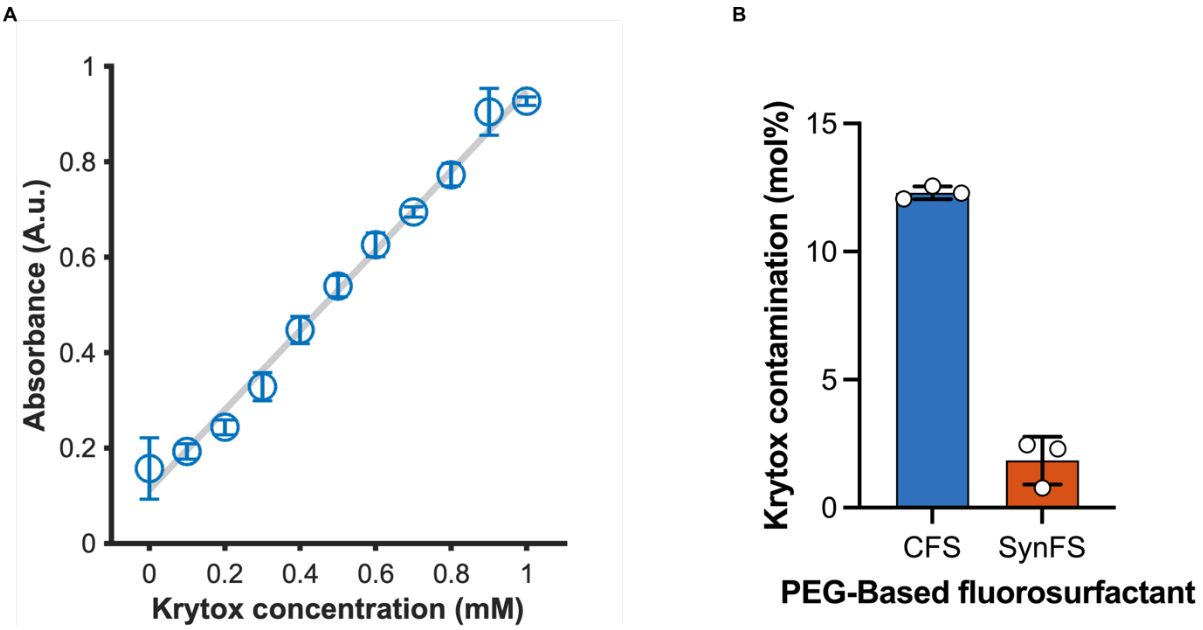


**Figure S8**: Krytox-mediated extraction of solutes. (**A**) Calibration curve of rhodamine 6G measured in the fluorous phase (HFE-7500) by absorbance spectroscopy at 530 nm ($R^2$ = 0.9912) at various concentration of Krytox supplemented within the fluorous phase. Mean ± S.D. (*n* = *3*) are presented. (**B**) Evaluation of the Krytox contamination within PEG-based fluorosurfactants by macroscopic partitioning Experiment. Aqueous solution of rhodamine 6G (1 mM) are exposed to the fluorous phase supplemented with different PEG-based fluorosurfactants: Commercially available PEG-based fluorosurfactant (CFS, 1.4 wt/wt% ≈ 3.8 mM) from Ran Technology, or a self-synthetized PEG-based fluorosurfactant (SynFS, 2.5 mM) for the present study. The rhodamine 6G signal was assessed into the fluorous phase by absorbance at 530 nm, and directly correlated to the Krytox impurity in the PEG-based fluorosurfactant due to a stochiometric association from (**A**).[13, 14] Mean ± S.D. (*n* = *3*) are presented.

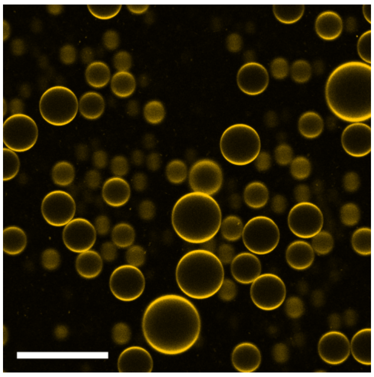

**Figure S9**: pH mediated assembly of droplet stabilized giant unilameller vesicles (dsGUVs). The aqueous phase was composed of 1.5 mM pH sensitive SUVs composed of DOBAQ/DOPG/DOPE/DOPC/Liss Rhod B labelled-DOPE (30/20/10/39.5/0.5 mol%) in 10 mM $KH_2PO_4$/$K_2HPO_4$ pH 7.4. The oil-surfactant mixture was composed of 2.5 mM PEG-based fluorosurfactant, and 10 mM Krytox in HFE-7500. The dsGUVs were produced by the one-pot shaking method, with an aqueous:oil ratio of 1:2 (50 : 100 µL). Scale bar: 50 µm.

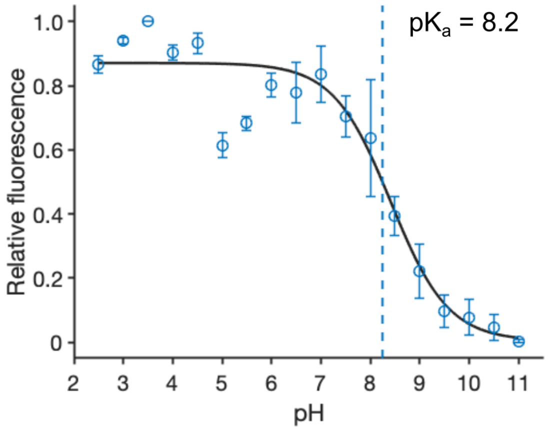


**Figure S10**: TNS assay to assess the apparent pKa of the pH sensitive lipid DODMA incorporated in SUVs composed of DODMA/DOPC (30/70 mol%). The increase in TNS signal correlated to the protonation of the amine functional group leading to an increase electrostatic attraction with TNS leading to an unquenching of its fluorescence. The pKa is defined as the half-maximum, corresponding to 50 percent of protonated DODMA lipid, extracted from a fitted sigmoid curve. The pKa is highlighted by the hatched vertical line. Mean ± S.D. are presented, *n* = *3*.

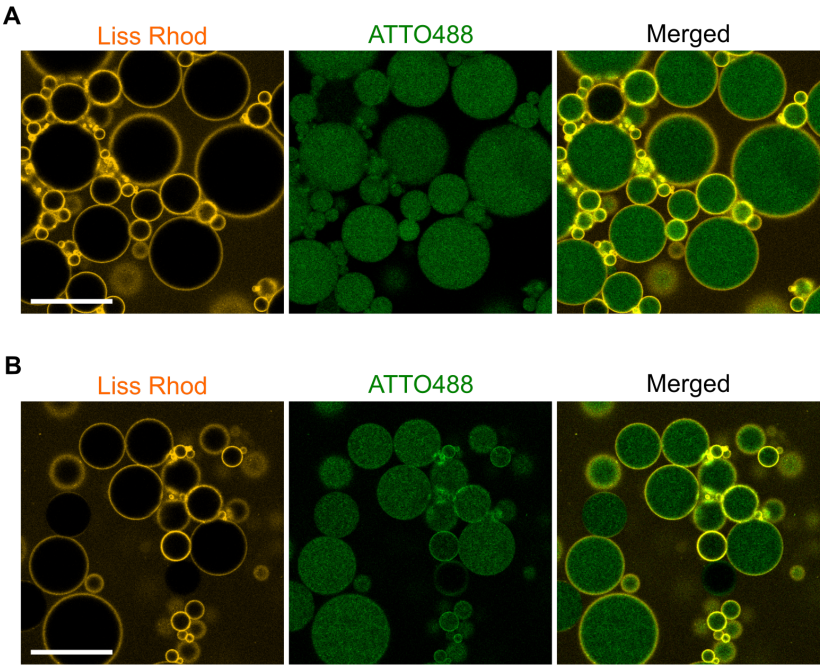


**Figure S11:** Self-assembly of multicompartment dsGUVs by Krytox acidification of the droplet lumen through co-encapsulation of two SUVs population. (**A**) Encapsulation of negatively charged SUVs composed of DOPG/DOPC/ATTO488-labelled DOPE (30/69.5/0.5 mol%) in 10 mM $KH_2PO_4/K_2HPO_4$, 140 mM KCl, pH of 7.4. (**B**) Encapsulation of neutral SUVs composed of DOPC/ATTO488-labelled DOPE (99.5/0.5 mol%) in 10 mM $KH_2PO_4/K_2HPO_4$, 140 mM KCl, pH of 7.4. In both cases, 1.5 mM of the pH sensitive SUVs composed of DODMA/DOPG/DOPC/DMG-PEG/Liss Rhod B-labelled DOPE (30/15/50.5/4/0.5 mol%) in 10 mM $KH_2PO_4/K_2HPO_4$, 140 mM KCl, pH of 7.4. The oil phase was composed of 2.5 mM PEG-based fluorosurfactant supplemented with 7.5 mM Krytox in HFE-7500.

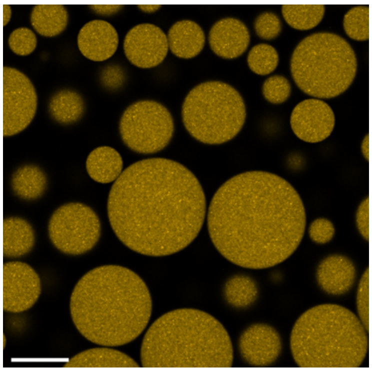

**Figure S12**: Production of dsGUVs in absence of $Mg^{2+}$ ions. Herein, 1.5 mM SUVs containing DOBAQ/DOPG/DOPC/DOPE/Liss Rhod B-labelled DOPE (30/20/39.5/10/0.5 mol%) in 30 mM Tris buffer pH 7.4 were encapsulated W/O droplets stabilized by 2.5 mM PEG-based fluorosurfactant and 10 mM Krytox in HFE-7500. The homogeneous fluorescence signal in the droplets' lumen highlight that no pH-mediated assembly is expected in these experimental conditions. Scale bar: 50 µm

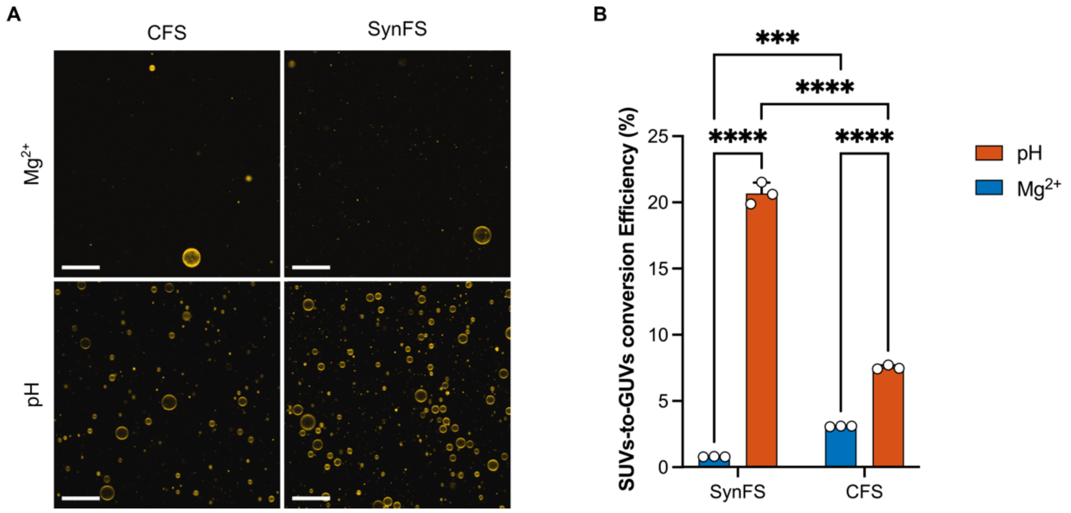


**Figure S13**: (**A**) CLSM images of the GUVs produced by pH- or $Mg^{2+}$-mediated assembly with the use of various PEG-based fluorosurfactant. In all cases, 1.5 mM of pH sensitive SUVs containing DOBAQ/DOPG/DOPC/DOPE/Liss Rhod B-labelled DOPE (30/20/39.5/10/0.5 mol%) were encapsulated into W/O droplets stabilized by the corresponding PEG-based fluorosurfactant. In the case of $Mg^{2+}$-mediated assembly, the aqueous phase was supplemented with 10 mM $MgCl_2$ and 30 mM Tris buffer pH 7.4. In the case of the pH-mediated assembly, the aqueous phase was 10 mM $KH_2PO_4/K_2HPO_4$, 140 mM KCl, pH 7.4. For the commercially available PEG-based fluorosurfactant (CFS), the oil-phase contained 1.4 wt/wt% CFS, and 10 mM Krytox in HFE-7500. In the case of SynFS, the oil-phase contained 2.5 mM SynFS, and 10 mM Krytox in HFE-7500. CLSM for the SynFS are presented in Fig. 4A, and included herein for visual comparison. (**B**) Two-way analysis of variance (ANOVA) of the SUVs-to-GUVs conversion efficiency as a function of the method of assembly, and the choice of the PEG-based fluorosurfactant assessed by fluorescence spectroscopy. Data for SynFS are presented in Fig 4B, but were included herein for the two-way ANOVA, and visual comparison. Mean ± S.D. ($n$ = 3) are presented. ***P<0.001; ****P<0.0001.

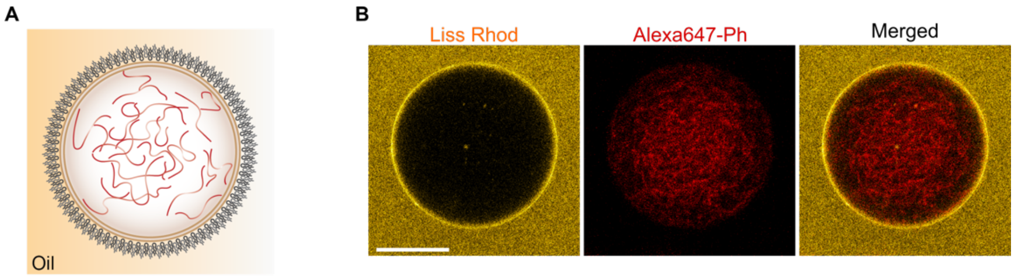


**Figure S14**: Reconstruction of an F-actin cytoskeleton within a synthetic cell by pH mediated assembly of droplet-stabilized GUV. (**A**) Scheme presenting the encapsulation of F-actin and pH sensitive SUVs within W/O droplets stabilized by PEG-based fluorosurfactant and Krytox. Herein, the pH sensitive SUVs fused to the droplet periphery to entrap the F-actin upon acidification of the droplet lumen by Krytox. The resulting synthetic cells were released in physiological condition leading to the production of minimal synthetic cell possessing an F-actin cytoskeleton as presented in figure 5B. (**B**) CLSM of droplet-stabilized GUVs encapsulating 1.5 mM pH sensitive SUVs composed of DOBAQ/DOPG/DOPE/DOPC/DMG-PEG/Liss Rhod B- labelled DOPE (30/20/10/35.5/4/0.5 mol%) and 5 µM of F-actin labelled with Phalloidin-Alexa647. Scale bar: 10 µm.

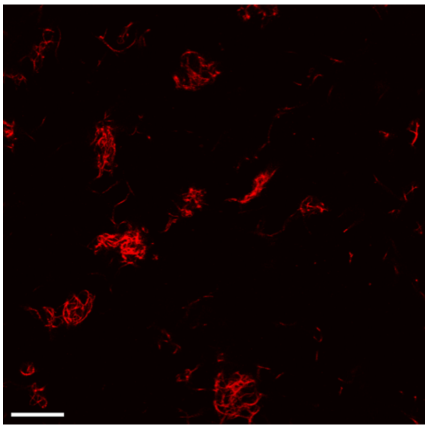

**Figure S15**: CLSM image of F-actin-Phalloidin-647 bundles and filaments adsorbed on the BSA-coated observation chamber. The F-actin located in the buffer following the release of the F-actin-loaded dsGUVs in physiological conditions originate from rupture of GUVs during the release process which liberate the entrapped F-actin into the buffer. Scale bar: 50 µm.

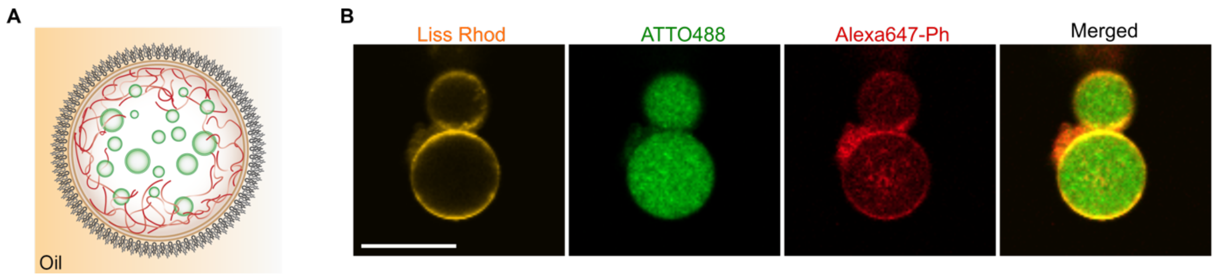


**Figure S16**: Reconstruction of an F-actin cytoskeleton and an endomembrane within a synthetic cell by pH mediated assembly of droplet-stabilized GUV. (**A**) Scheme presenting the encapsulation of both F-actin and negatively charged compartments along with pH sensitive SUVs within W/O droplets stabilized by PEG-based fluorosurfactant and Krytox. The pH sensitive SUVs fused to the droplet periphery to entrap the F-actin upon acidification of the droplet lumen by Krytox. The resulting synthetic cells were released in physiological condition leading to the production of minimal synthetic cell possessing both a F-actin cytoskeleton and and endomembrane system as presented in figure 5D. (**B**) CLSM of droplet-stabilized GUVs encapsulating 1.5 mM pH sensitive SUVs composed of DOBAQ/DOPG/DOPE/DOPC/DMG-PEG/Liss Rhod B-labelled DOPE (30/20/10/35.5/4/0.5 mol%), 1 mM of negatively charged SUVs composed of DOPG/DOPC/ATTO488-labelled DOPE (30/69.5/0.5 mol), and 5 µM of F-actin labelled with Phalloidin-Alexa647. Scale bar: 25 µm.

**Supplementary note 1**: Impact of the PEG-based fluorosurfactant on the SUVs to GUVs conversion for pH- and $Mg^{2+}$-mediated assembled dsGUVs.

To maximize the production of GUVs, the Krytox concentration within the oil-surfactant mix needs to be finely tuned, especially in the case of $Mg^{2+}$-mediated assembly of dsGUVs. Commercial PEG-based fluorosurfactants (CFSs) possess a none negligible contamination by Krytox,[14] which is the common source of PFPE monomer for the di- and triblock fluorosurfactant synthesis.[15-18] Also, the composition of CFSs correspond to an *a priori* variable mixture of di- and triblock copolymer revealed by MALDI-TOF analysis by Haller and collegues.[14] This potential variability results in frequent re-optimization of the experimental conditions to maintain an efficient GUV production. To overcome these variabilities and evaluate the impact of the PEG-based fluorosurfactant on the SUVs to GUVs conversion efficiency, we synthetized a PEG-based fluorosurfactant (SynFS) possessing minimal Krytox contamination, as showed by the partitioning assay (Fig S8). We applied click chemistry to generate the PEG-based fluorosurfactant, composed of $PFPE_{7000}$-$PEG_{1500}$-$PFPE_{7000}$. The click-based coupling was favored over classical amide bond formation due to its improved control, reproducibility of the synthesis[3] compared to traditional synthetic routes[15, 18, 19] and ease of purification steps, which minimized the potential Krytox contamination.

In the case of the pH-mediated assembly of GUVs, we observed a further improvement in both the absolute number of GUVs (Fig. S12A) and the SUVs to GUVs conversion efficiency (Fig. S12B) compared to $Mg^{2+}$ when the SynFS was used, but also compared to CFS. Interestingly, we noted that the conversion efficiency was reduced for $Mg^{2+}$ when the SynFS was used instead of the CFS. These results further highlight the necessity to optimize the experimental conditions for maximal release efficiency, and that *a priori* other interactions associated to the molecular structure of the surfactants would need to be accounted for. Nevertheless, the joint usage of SynFS and pH showed a significant improvement in SUVs to GUVs conversion efficiency to empower the high throughput production of multicompartment synthetic cells.

**Supplementary note 2**: Approximation of the time of diffusion of an SUV to the droplet periphery[7]

By assuming 3D Brownian motion, the mean square displacement, $\langle r^2(t) \rangle$, of a spherical particle correspond to

$$\langle r^2(t) \rangle = 6D\langle t \rangle \quad (1)$$

Where $D$ correspond to the diffusion coefficient of the particle. Thus, the average time $\langle t \rangle$ required for a particle to travel a distance of $\langle r^2(t) \rangle^{1/2}$ is

$$\langle t \rangle = \frac{\langle r^2(t) \rangle}{6D} \quad (2)$$

By applying the Stoke-Einstein equation of diffusion

$$D = \frac{k_B T}{6\pi\eta R} \quad (3)$$

Where $k_B$ is the constant of Boltzmann, $T$ the temperature, $\eta$ the viscosity of the solution and $R$ to the radius the particle in solution. Therefore, the average time $\langle t \rangle$ may be expressed as

$$\langle t \rangle = \frac{\langle r^2(t) \rangle}{6D} \times \frac{6\pi\eta R}{k_B T} = \frac{\langle r^2(t) \rangle \cdot \pi\eta R}{D k_B T} \quad (4)$$

By assuming a typical viscosity of $0.904 \times 10^{-3}$ Pa · s for a phosphate buffer saline (PBS), a temperature of 298 K, and particle radius of 50 nm, the time require by a SUVs travelling from the center of the W/O droplet possessing a mean diameter of 15 µm to reach the periphery ($\langle r^2(t) \rangle^{1/2}$ = 7.5 µm; $\langle r^2(t) \rangle$ = 56.25 µm$^2$) will be roughly 400 milliseconds.

Alternatively, an SUV of 100 nm diameter would have a $D$ of 24.13 µm$^2$ s approximated by the Stoke-Einstein equation (3).

**Supplementary note 3:** FRAP analysis

For the FRAP analysis, the normalized fluorescence intensity values were calculated as fellow:

$$I_{Normalized} = \frac{I_{Bleached} - I_{Reference}}{I_{Pre\ Bleached} - I_{Pre\ Reference}} \quad (5)$$

Where $I_{Bleached}$ correspond to the fluorescence intensity of the bleached spot, $I_{reference}$ is the fluorescence intensity on a unbleached/reference spot. $I_{Pre\ Bleached}$ and $I_{Pre\ Reference}$ were calculated by averaging the 10 measured fluorescence intensity values before the bleaching of the Bleached, and unbleached area, respectively. A non-linear least-square function was fitted to the normalized intensities from the recovery phase. The fit-function was:

$$f(t) = A(1 - \exp(-\lambda t)) + x_o \quad (6)$$

Where A and $\lambda$ are fit parameters, and $x_o$ correspond to the time point after bleaching, also referred as the start of the recovery phase. Then, by applying the protocol reported by Axelrod[20] and Soumpasis[21], the diffusion coefficient *D* of the lipids may be calculated via:

$$f = 0.32 \cdot \frac{r^2}{\tau_{1/2}} \quad (7)$$

Where $\tau$ is the half-recovery time, and *r* is the radius of the bleaching area.

**Supplementary note 4**: SUVs-to-GUV conversion efficiency calculation

The number of required lipid molecules to assemble a single giant unilamellar vesicle (GUV), $N_{Lip\ per\ GUV}$, can be written as:

$$N_{Lip\ per\ GUV} = 2 \cdot \frac{A_{SUV}}{A_{Head}} = 2 \cdot \frac{4\pi \cdot r_{GUV}^{2}}{A_{Head}} \quad (8)$$

Where $A_{SUV}$ is the area of a GUV possessing a radius $r_{GUV}$, and $A_{Head}$ is the area occupied by a single lipid head group.

The total number of lipids provided within the system, $N_{Lip}$, may be deduced from the total concentration of lipid used to produce dsGUVs:

$$N_{Lip} = c_{Lip} \cdot N_A \cdot V_{prod} \quad (9)$$

Where $c_{Lip}$ is the lipid concentration encapsulated during dsGUV formation, $N_A$ is the Avogadro number and $V_{prod}$ is the total volume of aqueous phase used to generate the W/O emulsion.

Therefore, the theoretical number of GUVs possible to produce with the provided lipids can be described by:

$$N_{GUVs\ Theo} = \frac{N_{Lip}}{N_{Lip\ per\ GUV}} = \frac{c_{Lip} \cdot N_A \cdot V_{prod} \cdot A_{Head}}{8\pi \cdot r_{GUV}^{2}} \quad (10)$$

Similarly, the experimental number of GUVs generated and released into physiological condition can be approximated by evaluating the lipid contain, and hence number of free-standing GUVs, of the released aqueous phase by:

$$N_{GUVs\ Exp} = \frac{N_{Lip\ free-Standing\ GUVs}}{N_{Lip\ per\ GUV}} = f \cdot \frac{c_{GUV} \cdot V_{Well} \cdot N_A \cdot A_{Head}}{8\pi \cdot r_{GUV}^{2}} \quad (11)$$

Where $f$ is a dilution factor associated to the dilution of the lipids, $c_{GUV}$ is the lipid concentration associated to the free-standing GUVs measured by a calibration curve generated with the precursor SUVs, and $V_{well}$ is the well's volume used to assess the fluorescence intensity by a plate reader measurement.

Then, the percentage of conversion efficiency, %Conversion, may be described as:

(12)

$$\%Conversion = 100\% \cdot \frac{N_{GUVs\ Exp}}{N_{GUVs\ Theo}} = 100\% \cdot f \cdot \frac{c_{GUV} \cdot V_{Well}}{c_{Lip} \cdot V_{prod}} \quad (13)$$

With an average $c_{GUV}$ of 7.75 µM, a well's volume of 100 µL, an initial lipid concentration $c_{Lip}$ of 1.5 mM lipids and an aqueous volume of production $V_{Prod}$ of 50 µL, the SUVs-toGUV conversion efficiency is roughly 20%.

**Note*** The factor *f* was evaluated as follow: Typically, 10 µL of free-standing GUVs were diluted to 100 µL (1:10 dilution), additionnaly, the initial 50 µL of lipid mixture used to assemble dsGUVs were released into a final 100 µL (thus, 1:2 dilution from initial lipid concentration). The resulting *f* factor corresponded to 20 in our experiment.

**Video S1**: pH sensitive SUVs entrapped in W/O droplets in presence of citrate buffer pH 5 using a two inlets microfluidics.

**Video S2**: Mechanical splitter module with a single inlet for production of multicompartment GUVs by microfluidics

**Video S3**: F-actin cytoskeleton inside a free standing GUV
